# TNF-NFkB-p53 axis restricts *in vivo* survival of hPSC-derived dopamine neuron

**DOI:** 10.1101/2023.03.29.534819

**Authors:** Tae Wan Kim, So Yeon Koo, Markus Riessland, Hyunwoo Cho, Fayzan Chaudhry, Benjamin Kolisnyk, Marco Vincenzo Russo, Nathalie Saurat, Sanjoy Mehta, Ralph Garippa, Doron Betel, Lorenz Studer

## Abstract

Ongoing, first-in-human clinical trials illustrate the feasibility and translational potential of human pluripotent stem cell (hPSC)-based cell therapies in Parkinson’s disease (PD). However, a major unresolved challenge in the field is the extensive cell death following transplantation with <10% of grafted dopamine neurons surviving. Here, we performed a pooled CRISPR/Cas9 screen to enhance survival of postmitotic dopamine neurons *in vivo*. We identified p53-mediated apoptotic cell death as major contributor to dopamine neuron loss and uncovered a causal link of TNFa-NFκB signaling in limiting cell survival. As a translationally applicable strategy to purify postmitotic dopamine neurons, we performed a cell surface marker screen that enabled purification without the need for genetic reporters. Combining cell sorting with adalimumab pretreatment, a clinically approved and widely used TNFa inhibitor, enabled efficient engraftment of postmitotic dopamine neurons leading to extensive re-innervation and functional recovery in a preclinical PD mouse model. Thus, transient TNFa inhibition presents a clinically relevant strategy to enhance survival and enable engraftment of postmitotic human PSC-derived dopamine neurons in PD.

**Highlights:**

- *In vivo* CRISPR-Cas9 screen identifies p53 limiting survival of grafted human dopamine neurons.
- TNFα-NFκB pathway mediates p53-dependent human dopamine neuron death
- Cell surface marker screen to enrich human dopamine neurons for translational use.
- FDA approved TNF-alpha inhibitor rescues *in vivo* dopamine neuron survival with *in vivo* function.

## Introduction

Parkinson’s disease (PD) is an increasingly common neurodegenerative disorder affecting around 10 million patients worldwide. PD causes an enormous clinical and financial burden to the affected individuals and to greater society (Bloem et al., 2021). The rapid increase in the number of PD cases globally has been referred to as a “Parkinson’s Pandemic” (Dorsey et al., 2018) stressing the need for novel therapeutic interventions, given that there are currently no disease-modifying or restorative therapies available. The characteristic motor symptoms in PD are caused by the specific and irreversible loss of midbrain dopamine neurons (Poewe et al., 2017; Surmeier et al., 2017). Therefore, cell therapy has been proposed as a novel therapeutic modality with the potential to achieve circuit level restoration of dopaminergic function. Proof-of-concept studies using human fetal midbrain tissue have shown promise with long-term improvement of motor-related symptoms in a subset of PD patients (Kefalopoulou et al., 2014; Li et al., 2016). However, access to human fetal tissue is limited and raises considerable logistical and ethical issues. Furthermore, grafting of fetal dopamine neurons showed high variability in clinical outcomes with two placebo-controlled trials failing to reach their primary endpoints (Freed et al., 2001; Olanow et al., 2003). Given these challenges using fetal tissue, human pluripotent stem cell (hPSC)-derived dopamine neurons have emerged as an alternative, more accessible, scalable and defined source of dopamine neurons that is the intense focus of ongoing, cell therapy efforts in PD (Barker et al., 2017; Kikuchi et al., 2017; Kim et al., 2021; Parmar et al., 2020; Schweitzer et al., 2020).

An unresolved challenge in dopamine neuron grafting is the limited *in vivo* cell survival. This is a critical issue given the need for sufficient dopamine neuron engraftment to achieve clinical improvement. Furthermore, low and variable survival rates (0.5% - 10%) likely contribute to inconsistent clinical outcomes and pose a risk for either under-or over-dosing patients in future hPSC-based trials. Finally, the need for injecting large numbers of cells to overcome limited *in vivo* survival poses the risk of triggering host inflammatory responses in response to extensive cell death of the grafted cells (Duan et al., 1995; Kriks et al., 2011; Tao et al., 2021; Winkler et al., 2005). Limited survival of target cell populations is a general problem for many hPSC-based cell therapy efforts including skeletal or cardiac muscle repair among others (Chong et al., 2014; Sun et al., 2022). During PD disease progression, certain mDA neuron subtypes are particularly susceptible such as substantia nigra (A9) neurons that undergo more rapid and extensive cell loss (Chen et al., 2021; Kamath et al., 2022; Liu et al., 2014; Panman et al., 2014; Pereira Luppi et al., 2021; Tao et al., 2021). Such disease-related vulnerability may translate into selective vulnerability during dopamine neuron grafting and thereby further limit the potency of dopamine neuron grafts, as A9 mDA neuron engraftment is critical for motor recovery. Understanding intrinsic factors that contribute to dopamine neuron vulnerability could enable improved grafting paradigms and offer insights into molecular determinants of PD disease vulnerability and progression.

Another challenge for developing hPSC-based cell therapies is the elimination of contaminating cell types such as non-dopaminergic neurons or non-neuronal lineages. For example, fetal tissue transplantation in PD patients caused graft-induced dyskinesia (GID) in a significant proportion of patients. It has been suggested that GID may be caused by contaminating serotonergic neurons present in fetal grafts (Politis et al., 2010). Furthermore, recent preclinical studies using with hPSC-derived cells pointed to the presence of other contaminants, such as TTR+ positive choroid plexus epithelial cells (Doi et al., 2020) or perivascular fibroblast-like populations (Tiklova et al., 2020). Surface marker sorting strategies have been proposed and implemented to enrich floor-plate intermediates or later stage dopamine precursors for transplantation (Doi et al., 2014; Lehnen et al., 2017; Samata et al., 2016), but none of these strategies specifically isolates early postmitotic dopamine neurons. First-in-human clinical trials using hPSC-derived lineages are based on grafting dopamine neuron precursors, rather than postmitotic dopamine neurons (Doi et al., 2020; Kikuchi et al., 2017; Kim et al., 2020; Kim et al., 2021; Kirkeby et al., 2017; Piao et al., 2021; Schweitzer et al., 2020). While this approach may circumvent the problem of selective vulnerability observed in A9 type dopamine neurons, use of early precursors rather than postmitotic neurons entails an increased risk for potential side-effects from “off-target” cell populations or from maintaining proliferation-competent precursors. It also limits the possibility of treating patients selectively with A9-, dopamine neuron subtype-specific grafts. Therefore, efficient engraftment of homogenous, post-mitotic dopamine neurons remains an important translational goal to further minimize safety-related concerns, and to reduce variability and optimize functional potency of hPSC-based grafts.

Pooled genetic screening using CRISPR/Cas9 technology has increasingly become the technology of choice to uncover causal genes driving specific phenotypes (Shalem et al., 2015). This approach has been widely adopted for *in vitro* studies, but has also been used to discover essential genes for *in vivo* tumor growth and metastasis (Chen et al., 2015), modulators for macrophage infiltration, and cancer immunotherapy targets (Wang et al., 2021b) as well as *in vivo* colon tumor suppressors (Michels et al., 2020). Here, we set out to systematically identify candidate mechanisms driving dopamine neuron death upon transplantation. We used a genetic Nurr1::H2B-GFP+ hPSC reporter line marking early postmitotic dopamine neurons to avoid confounding parameters such as proliferation in assessing survival. We identified p53 as a key factor restricting postmitotic dopamine neuron survival *in vivo* and described the temporal kinetics of p53 induction, host response, and mDA neuron death following transplantation. Transcriptomic analysis revealed TNF alpha (TNFa)-mediated activation of NFκB as upstream regulators of p53 induction, and we demonstrate that blocking TNFa-mediated signaling can rescue dopamine neuron death. To exploit those insights towards translational use, we identified a set of two cell surface markers to reliably purify NURR1::H2B-GFP+ post-mitotic dopamine neurons and thereby avoiding the need for a genetic reporter. Furthermore, we demonstrate that transient treatment of cells at grafting with an FDA-approved monoclonal antibody blocking TNFa (adalimumab) can dramatically improve postmitotic dopamine neuron survival *in vivo*, including ALDH1A1+ A9 type dopamine neurons, enabling efficient motor recovery in a preclinical PD mouse model. Our work addresses the mechanism underlying postmitotic dopamine neuron death at transplantation and establishes a clinically relevant strategy for cell-based therapies in PD patients.

## Results

### *In vivo* CRISPR/Cas9 screen identifies genetic drivers of postmitotic dopamine neuron death

We set out to develop an *in vivo* CRISPR-Cas9 screen to systematically address intrinsic factors that restrict the survival of hPSC derived post-mitotic dopamine neurons. Previously, we demonstrated that NURR1 (*NR4A2*) can serve as a reliable marker to denote early postmitotic dopamine neurons derived from hPSCs under the floor-plate differentiation paradigm (Chen et al., 2021; Riessland et al., 2019). FACS-based purification of *NURR1::GFP* positive cells yields nearly pure dopamine neurons populations in culture (Riessland et al., 2019) (**Figure S1A**), and gives rise to a highly homogenous and dense dopamine neuron graft *in vivo,* marked by TH expression (**Figure S1B**). However, grafting purified *NURR1::GFP+* dopamine neurons results in relatively poor survival (3%) in agreement with data from primary fetal dopamine neuron grafts. To screen for factors that limit *in vivo* survival, the *NURR1::H2B-GFP* reporter hPSC was further engineered to allow for doxycycline inducible Cas9 (iCas9) expression. This was achieved by TALEN-mediated targeting the AAVS1 safe harbor locus (Gonzalez et al., 2014) resulting in the iCas9/*NURR1::H2B-GFP* hPSC line (**Figure 1A**). To assess the efficiency of the iCas9 system for gene knock-out in our culture system, we stably transduced the iCas9/*NURR1::H2B-GFP* hPSC line which expresses Tomato driven by an EFS promotor (Addgene#89395) with an sgRNA targeting tdTomato (**Supplemental Table S2**). Lentivirus transduction was aimed at multiplicity of infection of 0.35 and stable Tomato expressing hPSCs were purified using FACS (**Figure S1C**). Upon doxycycline treatment from day 16 to day 25 of dopamine neuron differentiation (Kim et al., 2021), we observed efficient ablation of the Tomato signal (**Figure S1D**) without disrupting dopamine neuron induction based on the expression of the midbrain dopamine neuron markers NURR1 and FOXA2.

**Figure 1.**
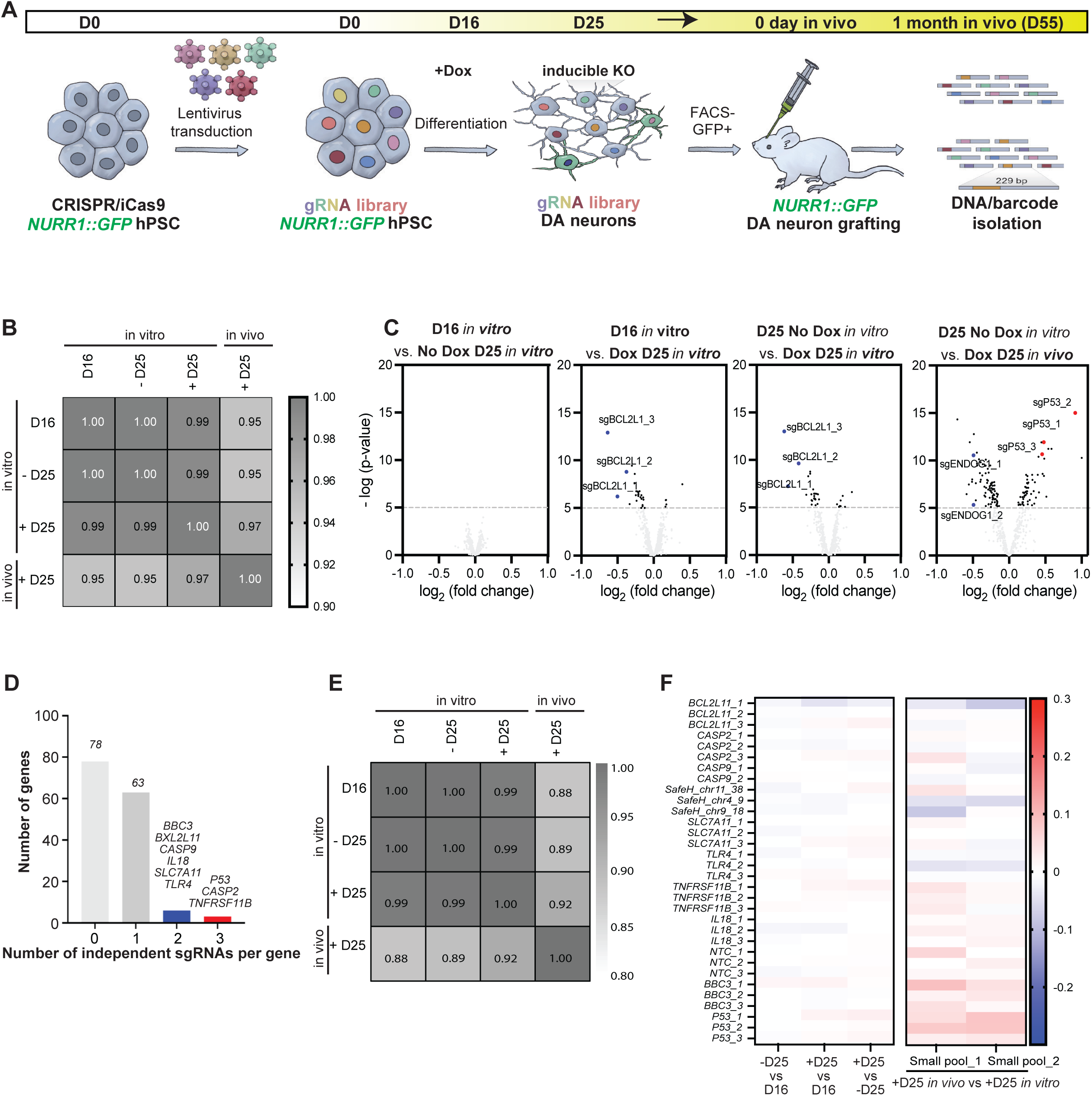
*in vivo* CRISPR/Cas9 screen identifies p53 as a limiting factor for the *in vivo* survival of hPSC-derived postmitotic dopamine neurons. (**A**) Schematic illustration of the pooled CRISPR/Cas9 screen (**B**) Pearson correlation abundance matrix of all guide RNAs across the different experimental conditions [day 16 *in vitro* (D16) vs. day 25 *in vitro* with no dox (-D25) vs. day 25 *in vitro* with dox (+D25) vs. day 25 *in vivo* with dox (D25)]. Scale bar range is from 0.9 to 1. 1 is the most correlated value. (**C**) Volcano plots comparing each experimental condition. Depleted sgRNAs are labeled in blue and enriched sgRNAs in red. (**D**) Enriched sgRNAs for *in vivo* grafted cells versus day25 *in vitro* cells, both treated with dox. Blue bar displays two sgRNAs and red bar shows three sgRNAs targeting for an indicated gene enriched in grafted cells than *in vitro* cells (see *criteria for sgRNA selection in the manuscript*). (**E**) Pearson correlation matrix plot for the pooled validation screen (library#2). Scale bar range is from 0.8 to 1. (**F**) Heatmap of all guide RNAs from pooled library#2 screen comparing day 16 (D16) vs. day 25 neurons without dox treatment (-D25) vs. day 25 with dox treatment (+D25) in culture (left). Heatmap of the same set of sgRNAs comparing +D25 *in vivo* vs +D25 in culture from two independent replicate screens (right). Red versus blue scale indicates enrichment of each sgRNAs in the surviving dopamine neurons in *in vivo* graft (+D25 *in vivo*) versus day25 cultured cells with dox (+D25 *in vitro*).

We next stably introduced a pooled, custom-designed, lentiviral library of 550 sgRNAs into the iCas9/*NURR1::H2B-GFP* hPSC line targeting a total of 150 genes (3 sgRNAs per gene) related to cell death pathways, such as apoptosis, necroptosis, pyroptosis, ferroptosis, and autophagy. The library further included 50 non-targeting and 50 safe harbor targeting control guides (**Supplemental Table S1 and S2**). Pooled lentiviral screening was performed at a multiplicity of infection of 0.35 (**Figure S1C**) to achieve single copy sgRNA integration per cell resulting in iCas9/*NURR1::H2B-GFP/*library hPSCs. Next, we directed the differentiation of the library-carrying hPSCs into dopamine neurons using our recently established protocol (Kim et al., 2021; Piao et al., 2021). At day 25 of differentiation, dopamine neurons were FACS-purified for co-expression of NURR1-GFP and gRNA-Tomato, and 800,000 postmitotic dopamine neurons were grafted into the striatum bilaterally into eight NOD/SCID IL2Rgnull (NSG) mice (**Figure 1A**). In parallel, we confirmed that FACS purified neurons gave rise to highly homogenous dopamine neuron *in vitro*, regardless of doxycycline treatment (**Figure S1E**). The use of NURR1+ neurons allowed us to examine cell survival of postmitotic dopamine neuron *in vivo* in the absence of any proliferating cells that could confound analysis. After 1 month post transplantation, the implanted cells were re-isolated by dissecting the graft based on fluorescence expression from thick tissue slices, followed by genomic DNA extraction (**Figure S1F**). We used a human specific PCR reaction detecting a human *PTGER2* gene to confirm the presence of the human cells within the xenograft sample (**Figure S1F**). Genomic DNA from both *in vitro* cultured and *in vivo* grafted cells were sequenced and showed robust sgRNA barcode library representation independent of Dox treatment (**Figure S1G**).

To identify the CRISPR targets that are specific to the survival of grafted dopamine neurons in contrast to those associated with dopamine differentiation, we assessed sgRNA ratios across both *in vitro* and *in vivo* samples (*in vitro* cultured cells: day 16, day 25 no dox, day 25 with dox, versus *in vivo* grafted cells: 1 month post grafting from day 25 with dox differentiation). The corresponding correlation matrix showed comparable sgRNA incorporation ratios regardless of dox treatment and the *in vitro* cell differentiation stage, though we observed a clear separation of the samples sequenced from the grafted group (**Figure 1B**). Analysis of all sgRNA ratios showed that, without dox treatment, we did not detect any enriched or depleted sgRNAs. However, in the dox treatment group, all 3 sgRNAs for BCL2L (BCLXL) were significantly depleted during *in vitro* differentiation (**Figure 1C**), demonstrating that BCLXL is a gene required for dopamine neuron induction in our culture system, consistent with the reduced numbers of dopamine neurons observed in Bcl-xl KO mice and overexpression data in neural stem cells or mouse ES cells showing increased dopamine neuron yield (Courtois et al., 2010; Savitt et al., 2005; Shim et al., 2004). Most importantly, we identified multiple significantly enriched sgRNAs in grafted dopamine neurons compared to the matched cells prior to grafting. Hits were identified based on two criteria: 1) sgRNAs enrichment above threshold [fold change > 0.3 (log_2_ value), and p value < 0.05]; 2) multiple independent gRNAs targeting the same gene identified as hits [fold change; more than 0.25 (log_2_ value), and p value; p < 0.05]. Using these as criteria, we identified 9 genes as hits restricting *in vivo* dopamine neurons survival and highlighting apoptosis and inflammation as the main pathways involved (**Figures 1C and 1D**). In contrast, we found that only ENDOG gene (2 sgRNAs) is significantly depleted in grafted *vs* culture dopamine neuron.

To further validate the 9 genes identified as hits in the screen, we generated stable hPSCs transduced with 33 sgRNAs targeting those 9 genes (3 sgRNAs per a gene) in addition to 6 non-targeting and safe harbor control guides (**Supplemental Table S2**) resulting in an hPSC line termed iCas9/*NURR1::H2B-GFP/*library#2. The overall representation of sgRNAs was comparable across all conditions (**Figure S1H**), and sgRNA incorporation ratios were again most distinct in the grafted group (**Figure 1E**). The two independent replicate screens using the small pool validation sgRNA library (iCas9/*NURR1::H2B-GFP/*library#2 hPSCs) identified p53 as the most enriched hit (**Figure 1F**) and indicating that postmitotic dopamine neuron death upon transplantation is likely driven in a p53-dependent manner.

### p53 knockout (KO) results in improved dopamine neuron survival

To further examine the role of p53 in dopamine neuron survival following transplantation, we generated an iCas9/*NURR1::H2B-GFP* hPSC line carrying a stable p53-targeting sgRNA. While the p53 pathway has been previously implicated in the survival of fetal ventral-mesencephalic graft in rats (Chou et al., 2011), the p53 effect on the survival of postmitotic human dopamine neurons *in vivo* remains unknown. Furthermore, there is no information on the kinetics of p53-mediated dopamine neuron death post grafting. We differentiated the p53 iCas9/*NURR1::H2B-GFP* hPSC line into dopamine neurons followed by FACS-based purification for NURR1::H2B-GFP and sgRNA::Tomato. There was no improved *in vitro* dopamine neuron survival or proliferation in the dox treated (p53 KO) group, with both dox-treated and non-treated cultures yielding highly enriched dopamine neurons expressing TH and FOXA2 at 2 weeks post-sorting (**Figure S2A**). We grafted dox treated (p53 KO) versus non-treated (isogenic p53 WT) dopamine neurons into the left versus right striatum of the same mouse, to minimize variability that could be related to grafting independent animals (**Figure 2A**). At 1 month post transplantation, we determined overall graft composition and size using immunofluorescence followed by stereological quantification (**Figures 2B and C**, **Figures S2B–D**). All the surviving p53 KO dopamine neurons showed expression of floor-plate and dopamine neuron identity markers such as FOXA2, TH and NURR1-GFP signal (**Figures S2B–D**). We did not find evidence of KI67 expression, which marks proliferating cells, in p53 KO or in isogenic control grafts (**Figure S2E**). We used stereological methods to determine graft volume and the total number of surviving *NURR1::GFP* positive neurons per 100,000 grafted cells. We found that p53 KO grafts contained 13,667 (mean) ± 3,590 (S.E.M) *NURR1::GFP* dopamine neurons compared to 2,955 (mean) ± 435 (S.E.M) in wild-type neurons (p = 0.0087, paired t-test, n = 5) (**Figure 2C**). The volume of the p53 KO graft was 0.088 mm^3^ (mean) ± 0.020 (S.E.M) mm^3^ while the wild-type graft was 0.018 mm^3^ (mean) ± 0.003 mm^3^ (S.E.M) mm^3^ per 100,000 grafted cells (p = 0.0321, paired t-test, n = 5) (**Figure 2C**). In addition to the increased volume and increased numbers of NURR1+ neurons, p53 KO grafts also showed an increased proportion of ALDH1A1+ A9-type dopamine neurons. In contrast, the proportion of CALB1 positive (A10) neuron was comparable between p53 KO and p53 WT grafts (**Figures 2D–G**), suggesting a particular vulnerability of A9 but not A10 dopamine neuron subtype upon grafting, which is alleviated in p53 KO grafts.

**Figure 2.**
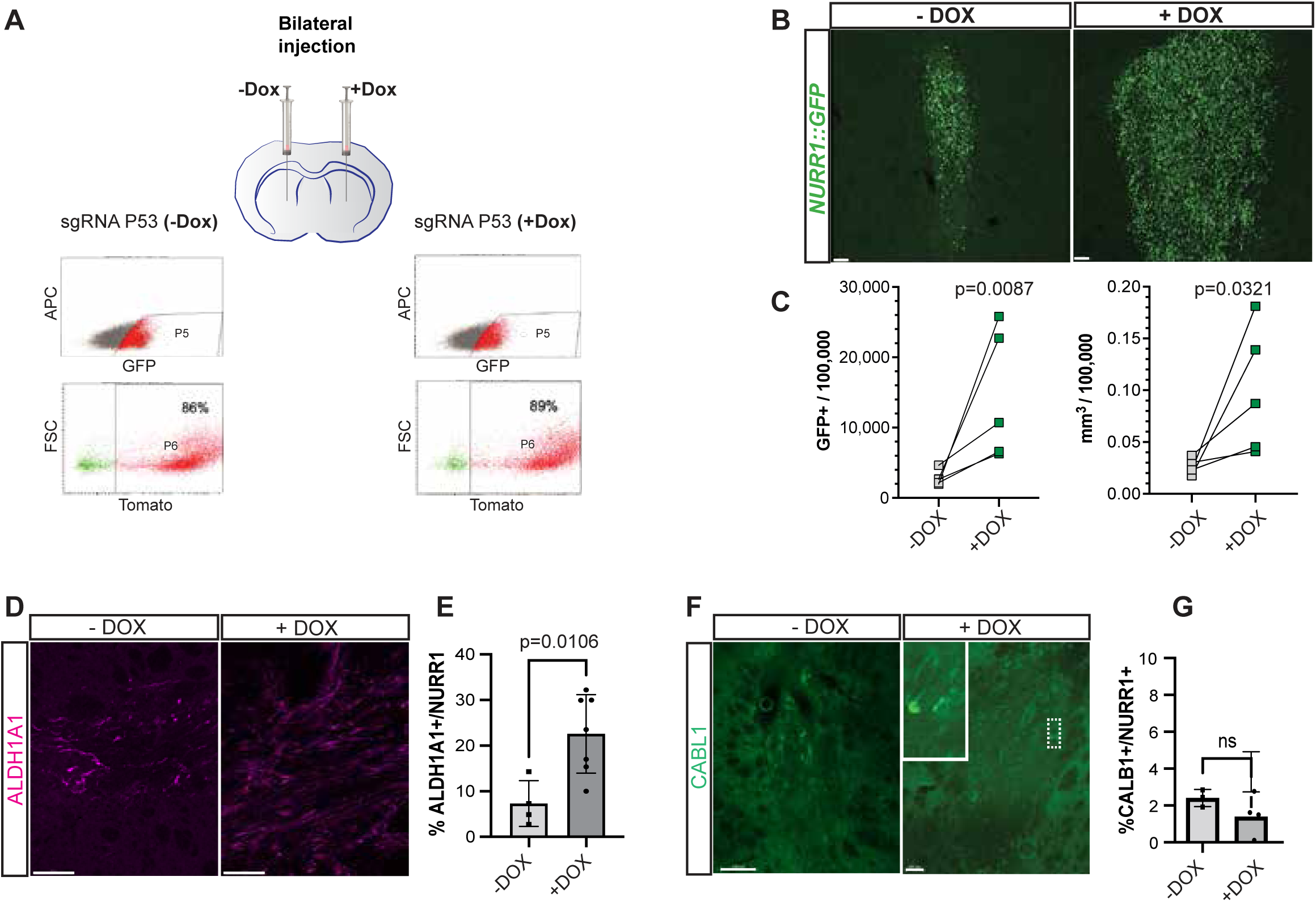
Characterization of p53-induced dopamine neuron death during transplantation. (**A**) FACS strategy for injecting enriched postmitotic dopamine neurons expressing NURR1::GFP and sgRNA-P53-tdTomato. Each dot in the scatterplot indicates a single dopamine neuron at 25 DIV. Dox treated from day16 to 25 (+DOX, P53 knock-out; KO) or non-treated (-DOX, isogenic P53 wild-type; WT) dopamine neurons are isolated by FACS at day25 based on NURR1::GFP signals (upper panel: P5), followed by sgRNA::tdTomato signal (bottom panel: P6). GFP+/tdTomato+ dopamine neurons (P6 population: -DOX vs +DOX) are bilaterally injected into the striatum of adult NSG mice. (**B, C**) Representative confocal images of dopamine neuron grafts at 1 month post transplantation stained with antibodies against GFP (equivalent to NURR1), scale bar = 100 μm. **(B)** and stereological analysis for the number (using optical fractionator, left) and volume (cavalier estimator, right) of the surviving dopamine neurons at 1 month post transplantation. n= 5 for each condition (**C**). (**D, F**) Representative confocal images of ALDH1A1 (A9 type dopamine neuron marker, D) and CALB1 (A10 type dopamine neuron marker, F). Scale bar = 100 μm. (**E, G**) Quantification of the percentage of A9 (ALDH1A1, E) and A10 (CALB1, G) dopamine neurons per NURR1 expressing dopamine neurons at 1 month post transplantation. See *Materials and Method.* *p<0.05 (paired t-test). ns. = not significant.

### Temporal analysis of p53-mediated dopamine neuron death and host response

Next, we sought to address the kinetics of the p53-dependent neuronal cell death following transplantation. We assessed three main stages as potential drivers of the cell death: 1) Mechanical damage such as shear forces during loading of the cells into the needle 2) Short-term survival factors immediately post transplantation 3) Longer-term effects in established graft. During cell loading and ejection we did not observe induction of p53, and downstream genes as measured by qRT-PCR (**Figure S2F**), consistent with our previous work showing no obvious signs of cell death after loading and ejecting cells from needle (Kim et al., 2021). We next tracked p53 and cleaved caspase 3 (CC3) induction by immunofluorescence at sequential time points post grafting and observed robust induction of both markers in about 30-40% of the dopamine neurons at 24 hours post-transplantation (dpt). Interestingly, at 4 hours, p53 expression in dopamine neurons was still very low (**Figures 3A and B, D and E**). At 3 dpt, the percentage of p53 positive cells was already greatly diminished compared to 1 dpt while the percentage of CC3 and TUNEL positive, apoptotic cells remained high at 3 dpt. These data indicate that p53-induced neurons at 1 dpt may trigger an apoptotic program executed over several days. In parallel, the rapid engagement of cell death results in a dramatic decrease in cell density of the graft comparing 1 dpt versus 3 or 7 dpt (**Figures 3A–C**). By 7 dpt and beyond, the wave of programmed cell death completely subsided. Confirmation of these findings by TUNEL assay, which probes for apoptotic DNA fragmentation, showed strong TUNEL labeling starting at 1 dpt, peaking at 3 dpt and with no remaining TUNEL positive cells by 7 dpt (**Figures 3C, F**).

**Figure 3.**
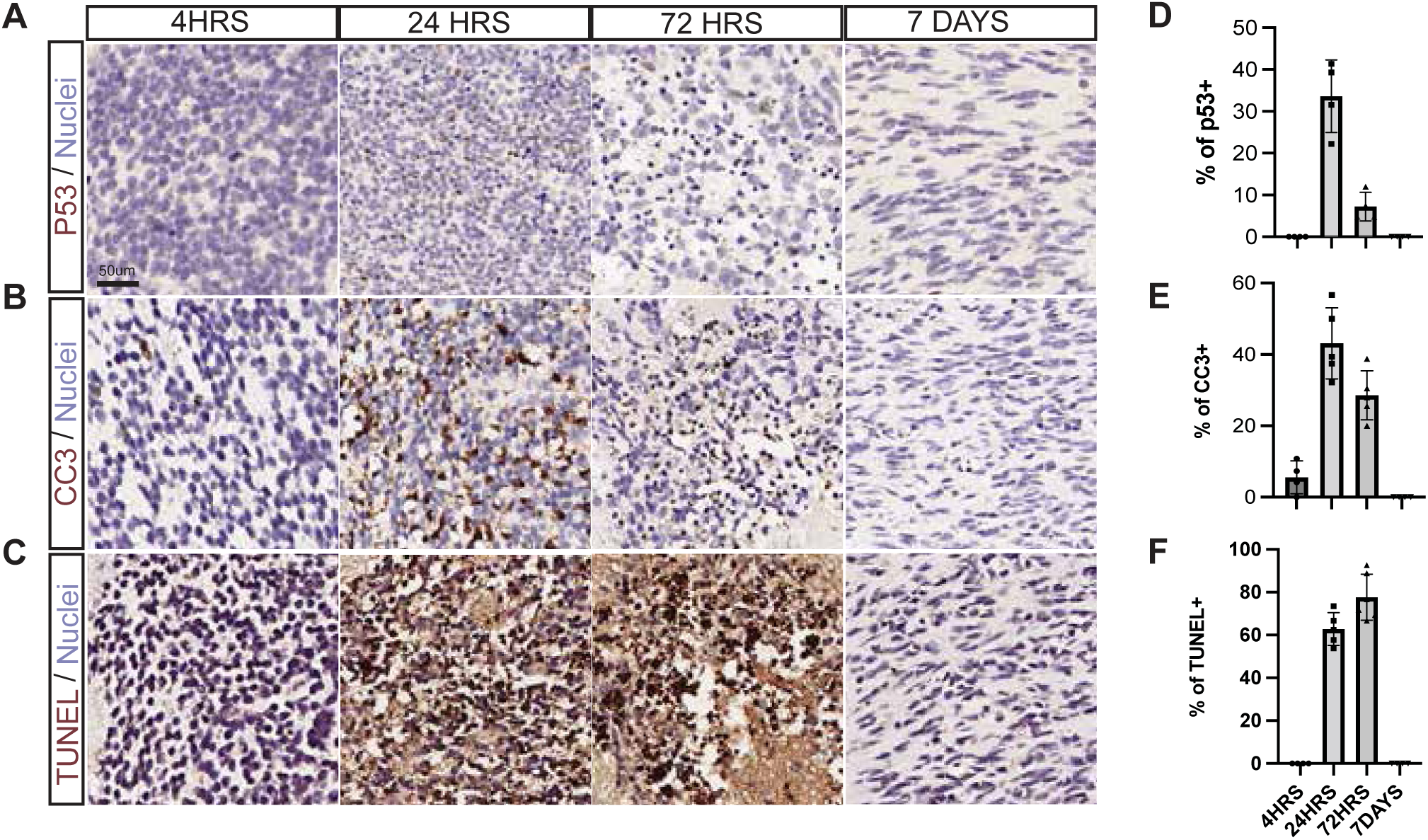
Temporal kinetics of the p53-mediated dopamine neuron death-related pathways post implantation. (**A-F**) Representative immunohistochemistry image (**A-C**) and quantification of the percentages (**D-F**) of the grafted dopamine neurons at different time points (4 hpt, 24 hpt, 72 hpt, 7 dpt) for p53, cleaved caspase 3 (CC3), and TUNEL as apoptosis markers in dopamine neuron grafts upon transplantation. All markers show robust induction at 24hr. Scale bar = 50 μm. n = 4 independent experiments.

Next, we characterized the dynamics of immune-related cells at the graft site to assess whether the host response precedes or follows p53-induced dopamine neuron death. We observed that microglia and astrocytes concurrently polarize their processes towards the transplants at 1 dpt, a time point when there is already ongoing dopamine neuron apoptosis. However, at 4 hours and at 1 dpt most glial cells were located outside the graft core, suggesting that they are responding to rather than driving the initial wave of dopamine neuron cell death (**Figures S3A**). At 3 dpt, the inflammatory response peaked with extensive polarization of glial cells, in particular the microglial population entering the graft core and encapsulating apoptotic cell bodies (**Figure S3A**, upper panels). In addition, we observed strong GFAP staining, marking activated astrocytes at the graft/host interface which persisted by 7 dpt creating an “astrocyte border” at the graft host interface (**Figure S3A**, lower panels). Finally, we observed evidence of vascular recruitment to the graft at 3 dpt by H&E staining (**Figure S3B**) and neutrophil invasion marked by Ly6G present prior to 24 hours post grafting (**Figures S3C**). Despite ongoing cell death, grafted surviving neurons began to extend axons as early as 1 dpt, illustrated by STEM121 positive fiber outgrowth, from the graft core (**Figure S3D**). These data suggest that *in vivo* survival of dopamine neurons (fiber outgrowth versus apoptotic death) is determined within the first 24-hour post engraftment. Overall, our data reveal a rapid p53-dependent cell death of hPSC-derived postmitotic dopamine neurons *in vivo* which triggers a host response peaking at 3 dpt followed by clearing of apoptotic nuclei and the subsiding of dopamine neuron death by 7 dpt.

### TNFa-NFκB pathway is an upstream regulator of p53-dependent apoptosis in engrafted dopamine neurons

Given the well-known role of p53 as a tumor suppressor, stable manipulation of this gene in hPSC-derived dopamine neurons is not suitable for translational applications. Furthermore, at the mechanistic level, it is important to understand the upstream molecular pathways that lead to p53 induction at 1 dpt. We performed bulk RNAseq analysis to compare gene expression of grafted NURR1+ neurons, re-isolated at 1 dpt from the mouse brain, versus that of the matched FACS-purified neurons isolated immediately prior to transplantation (d0) or analyzed at 1 day of *in vitro* culture. Principal component (PCA) and dendrogram analyses demonstrated that the grafted dopamine neurons exhibited the most distinct transcriptional pattern compared to either sorted or *in vitro* cultured dopamine neurons (**Figure 4A and Figure S4A**). Gene ontology analysis of the 279 differentially expressed genes (DEG) upregulated in 1 day *in vivo* grafted versus 1 day *in vitro* cultured neurons showed strong enrichment of TNFa signaling via NFκB signaling, apoptosis, hypoxia, and p53 pathways (**Figures 4B, C**) whereas the 374 DEG downregulated in the *in vivo* grafted vs. the cultured neuron were associated with apical junction, mTORC1 signaling, and cholesterol homeostasis (**Figure 4B and Figure S4B**). Gene Set Enrichment Analysis (GSEA) for TNFa signaling (**Figure 4D and Figure S4C**) and unbiased analysis of all enriched pathways in day 1 grafted neurons vs. day 1 *in vitro* plated neurons confirmed apoptosis, p53 and TNFa signaling among the most consistently enriched pathways (**Figure 4E**).

**Figure 4.**
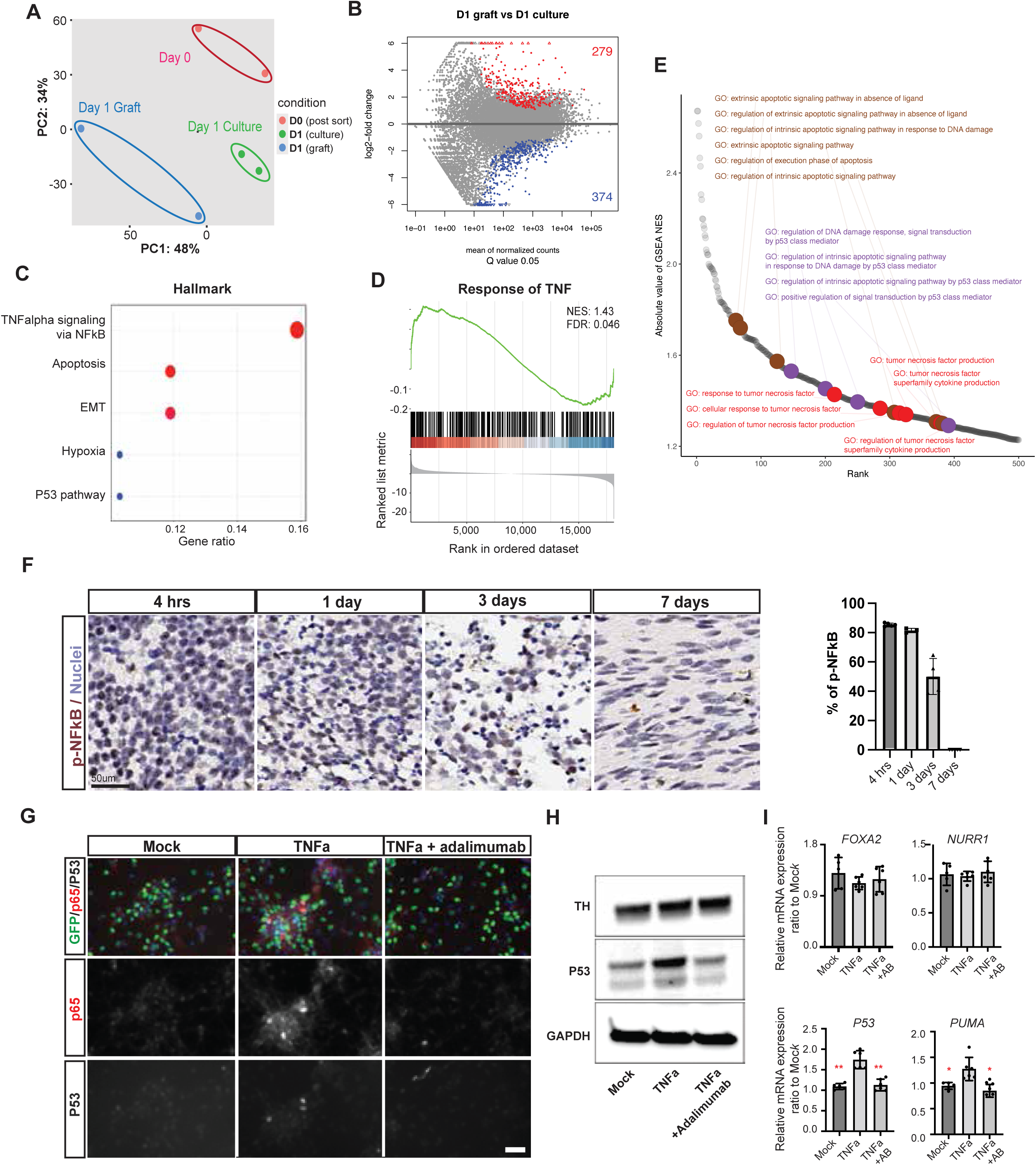
TNFa-NFκB pathway is an upstream trigger of p53-dependent dopamine neuron death in the graft. (**A**) PCA plot of bulk RNAseq data for sorted dopamine neurons either immediately post FACS (day 0, D0), *in vitro* cultured for 1 day post sorting (day 1 culture, D1 culture) or *in vivo* grafted for 1 day (day 1 graft, D1 graft). (**B**) Differentially expressed gene (DEG) analysis between D1 culture versus D1 graft. (**C**) Hallmark pathway analysis on the upregulated genes for functional categories in D1 grafted vs D1 cultured neuron. (**D**) Normalized enrichment score (NES) analysis of enriched TNF-related genes in D1 graft *vs*. D1 culture neuron. (**E**) Unbiased gene set enrichment analysis (GSEA) identifies Apoptosis, P53, and TNFa as the GO terms most frequently enriched in D1 graft versus D1 culture neuron. (**F**) Left panels: Representative immunohistochemistry images of phosphorylated NFkappaB (p-NFκB) in grafted neurons at distinct time points post transplantation, scale bar = 50 μm. Right panel: Quantification of the percentages of p-NFκB positive cells among total cells within the graft, n = 3 independent experiments. (**G**) Representative immunofluorescence images of NURR1::GFP sorted dopamine neurons *in vitro* for the induction of p53 and NFkB-p65 comparing mock *vs.* TNFa *vs.* TNFa and monoclonal antibody against TNFa (adalimumab), treated conditions for 1 day. (**H, I**). Western blot and qRT-PCR of gene expression profiles of the three groups listed in (**G**) for TH and P53 (**H**) and for the midbrain dopamine neuron markers FOXA2 and NURR1, P53, and PUMA downstream target of P53 (**I**). N > 3 independent experiments.

We next monitored activation of nuclear factor kappa B (pNFκB) signaling as a further readout of TNFa activation in the grafted cells. Time-course analysis by immunofluorescence showed strong induction of pNFκB in grafted neurons as early as 4 hours and 1 dpt. Nearly 80% of the grafted neurons showed pNFκB expression with a dramatic decrease in the percentage of pNFκB positive neurons by 3 dpt (**Figure 4F**) and beyond. In contrast, at 1 day post plating, nuclear NFκB expression was not induced in dopamine neuron replated *in vitro* (**Figure S4D**). These data suggest that TNFa-NFκB signaling cascade is induced shortly after dopamine neuron engraftment and likely prior to p53-dependent apoptosis, pointing to TNFa-NFκB pathway as a candidate upstream regulator.

To validate a causal link between TNFa-mediated NFκB signaling and p53 induction, we treated FACS-purified dopamine neurons with recombinant TNFa *in vitro*. Under control conditions there was no evidence of nuclear NFκB (based on p65 expression) or p53 expression (**Figure S4D**) further supporting the notion that this pathway is selectively triggered in grafted dopamine neurons *in vivo* but not upon replating *in vitro*. However, when treated with TNFa, cultured dopamine neurons showed induction of both nuclear p65 and p53, a response that could be blocked by co-treatment with the TNFa-blocking monoclonal antibody adalimumab (**Figures 4G and H**). Interestingly, only a subset of TNFa-treated dopamine neurons was positive for p53 but those all co-expressed NFκB-p65, further supporting a causal link. In addition, TNFa treatment led to a significant transcriptional activation of p53 and its downstream gene, PUMA that was abolished by co-treatment with adalimumab (**Figure 4I**).

### Single cell RNA sequencing of day 1 grafted neurons identifies molecular signature of surviving dopamine neurons and de-differentiation signature in apoptotic dopamine neurons

To further define the transcriptional landscape of grafted dopamine neurons, we performed single cell mRNA sequencing from p53 wild-type (WT) and p53 knock-out (KO) grafted neurons, re-isolated from the mouse brain at 1 dpt. Combined clustering of p53 WT and p53 KO cells showed highly overlapping distribution of clusters, implying that the cell state is not changed in dopamine neuron upon p53 KO (**Figures 5A and B, Figures S5A–C**). Based on *PBX1*, a marker particularly highly expressed in dopamine neurons (Villaescusa et al., 2016), and on the expression of the neuronal marker *MAP2*, >90% of the cells exhibited features of postmitotic, dopamine neuron identity with very low expression of the proliferation marker, *MKI67*, in both p53 WT and KO neurons (**Figures S5B**). By comparing our data to an available dataset from human fetal dopamine neuron development (La Manno et al., 2016), most of the cells isolated at 1dpt were annotated as neuroblasts, compatible with early NURR1 positive identity while a small fraction of the cells was annotated as floor plate progenitors (**Figure 5C**).

**Figure 5.**
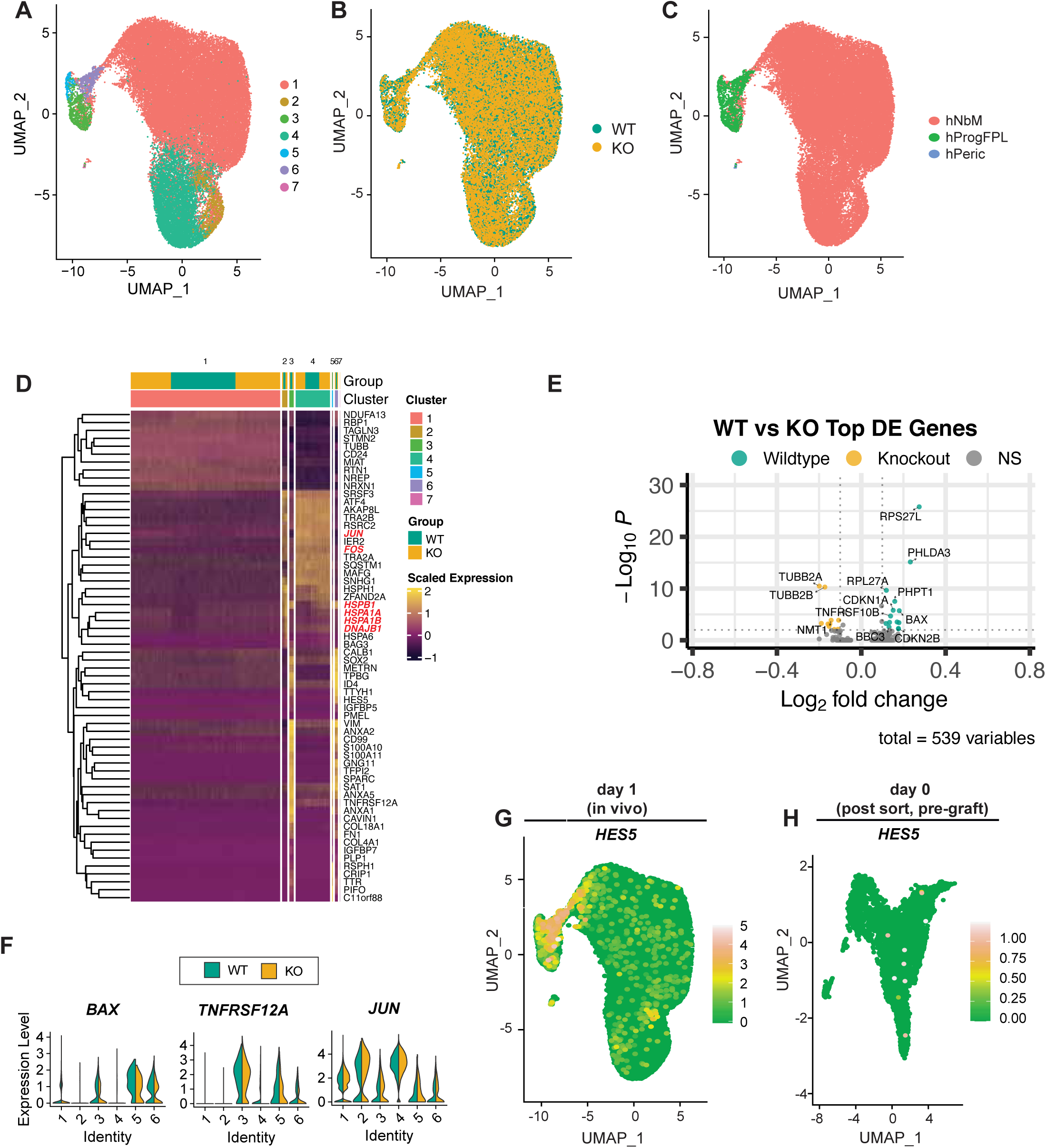
Single cell RNA sequencing of grafted neurons identifies JUN-related survival and dedifferentiation-associated cell death signatures following transplantation. (**A-C**) UMAP plot of scRNA-seq from P53 WT and KO grafted cells at 1 day post transplantation. Color coded by cell clusters (**A**), P53 WT and KO genotypes (**B**), and annotated cell types including neuroblasts (hNbM), floor-plate progenitor (hProgFPL), and a very small portion of pericytes (hPeric) (**C**). (**D**) Heatmap of top enriched gene-set in each cluster of P53 WT versus P53 KO cells at 1 day post transplantation, demonstrating highly increased cell death-related genes in clusters 3, 5, and 6 and survival related genes in clusters 2, 4. Red color indicates survival related genes. (**E**) Volcano plots of differentially expressed genes of P53 downstream genes, such as *BAX, CDK1NA, CDKN2B, BBC3 (PUMA)*, and *PHPT1*, are significantly increased in P53 WT versus P53 KO grafted dopamine neurons from clusters 3, 5, and 6. (**F**) Violin plots of BAX, TNFRSF12A, and JUN positive cells among the clusters. The cluster 7 is excluded due to a very small portion of cells. (**G, H**) HES5 positive cells, specifically to clusters 3, 5, and 6, mark de-differentiated signature in UMAP from 1day post graft (**G**) that is barely expressed in the sorted cells prior to grafting (**H**).

We next identified clusters that displayed molecular signatures of apoptosis or survival. Differential analysis of p53 WT versus p53 KO identified clusters 3, 5, and 6 with significantly increased expression of p53 downstream genes, such as *BAX, BBC3, CDKN1A*, and *PHPT1* (**Figures 5D–F**). In these clusters, TNFRSF12A was more robustly expressed in both p53WT and p53KO cells relative to the other clusters, supporting our findings that TNFa mediates a p53-dependent apoptosis pathway in the graft (**Figure 5F**). Interestingly, apoptosing clusters showed expression of HES5 and were annotated as floor plate progenitors. However, we did not observe any evidence of proliferation based on *MKI67* expression, suggesting that this cluster may reflect a dedifferentiation state rather than a bona fide, proliferating floor plate progenitor signature (**Figures 5G and H**). Furthermore, the progenitor signature was specific to immediately grafted neurons (1 dpt) and not observed in matched cells immediately pre-engraftment cells (**Figure 5H**). These data argue that the progenitor-like phenotype is induced in those clusters within 24 hours post-engraftment and likely does not represent a contamination from imperfect sorting.

Conversely, clusters 1, 2, and 4 show high expression of survival signature characterized by the expression of *ATF4, JUN, FOS, HSPA6, HSPA1A, HSPA1B*, and *DNAJB1* (**Figure 5D**), highlighting that hPSC-derived dopamine neurons are under high cellular stress during engraftment procedure such as due to axotomy, endoplasmic reticulum stress, or DNA damage. Differential analysis between p53 WT and KO identified the survival marker JUN as significantly upregulated upon p53 KO (**Figure 5F and Figure S5D**), emphasizing that blocking p53 expression imparts survival benefits to grafted dopamine neurons.

One major unresolved question is the source of TNFa ligand and the mechanisms responsible for triggering TNFa induction. Further study is needed to determine whether the initial increase in TNFa ligand expression is related to cell damage from the grafting procedure itself, and to what extent graft-versus host-derived TNFa is responsible for mediating the subsequent activation of a NFκB-dependent and p53-mediated apoptotic cell death response.

### Surface marker screen defines antibody strategy for purifying postmitotic dopamine neuron

To translate our findings from purified NURR1::H2B-GFP positive, postmitotic dopamine neurons into a platform suitable for future clinical translation, we sought to establish a surface marker-based sorting strategy obviating the need for a genetic reporter line. We performed a high-throughput flow-based cell surface marker screen using 385 validated antibodies (Princess Margaret Cancer Centre, University Health Network Antibody Core Facility) (Gedye et al., 2014) (**Supplemental Table 3**). We identified three CD markers that were anti-correlated with NURR1 levels (CD49e, CD99, CD340) and two markers positively correlated with NURR1 (CD184, CD171) when assessed in *NURR1::H2B-GFP* hPSC-derived dopamine neurons at day 25 (**Figure 6A and S6A**). After extensive characterization of sorted cells following either single and/or double CD marker enrichment, we found that CD49e-low/CD184-high expressing cell were most highly enriched for postmitotic dopamine neurons, based on GFP expression (**Figure 6B**). Similarly, NURR1 mRNA expression was significantly enriched in CD49e low / CD184 high cells compared to unsorted or single sorted or double sorted cells with CD49e-low and CD171-high (**Figure 6C**). We further confirmed the purity of postmitotic dopamine neurons *in vitro* by staining for FOXA2, NURR1::GFP, and TH at 2 weeks post sorting (**Figure 6D**). Moreover, we performed short-term intrastriatal grafting of CD49e-low/CD184-high sorted dopamine neurons, resulting in near homogenous TH+ neuron grafts devoid of MKI67 proliferating cells at 1 month post transplantation (**Figure 6E and Figure S6B**). These data demonstrate the feasibility of enriching for postmitotic dopamine neurons *via* a novel CD marker strategy suitable for *in vivo* transplantation.

**Figure 6.**
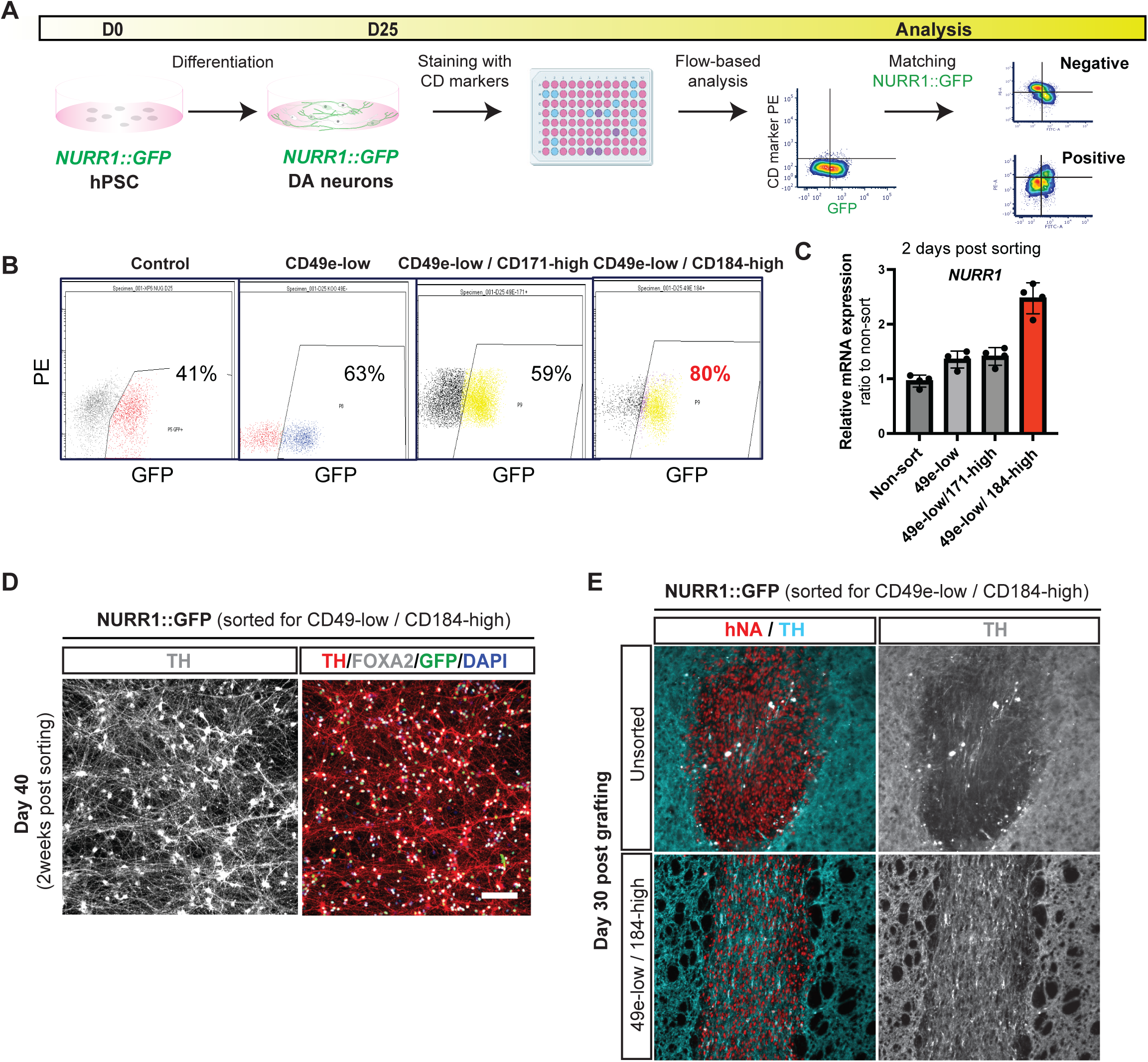
High-through flow-based cell surface marker screen identifies novel CD marker to purify NURR1 stage postmitotic dopamine neuron for translational use. (**A**) Schematic illustration of the flow-based CD marker screen to enrich for postmitotic dopamine neurons matching genetic NURR1::GFP reporter expression. (**B**) FACS plot of % NURR1::GFP populations corresponding to each sorting strategy (Control, CD49e-low, CD49-low/CD171-high, CD49e-low/CD184-high), indicating CD49e-low/CD184-high double CD marker sorting leads to the most enriched dopamine neuron population expressing NURR1::GFP. (**C**) Gene expression of NURR1 via qRT-PCR assay 2 days post sorting using each sorting strategy from (**B**). (**D**) Representative immunofluorescence image of CD49e-low/CD184-high double sorted dopamine neurons at day 40, giving rise to pure dopamine neuron culture co-expressing NURR1::GFP, FOXA2, and TH. Scale bar = 100 μm. (**E**) Short term *in vivo* histology analysis at 1 month post grafting of CD49e-low/CD184-high double sorted graft compared with unsorted cells. The CD49e-low/CD184-high sorted neuron grafts are composed of densely packed dopamine neurons in contrast to unsorted grafts which yield a lower percentage of dopamine neurons as detected by human nuclear antigen (hNA) and TH immunofluorescence-staining.

### Adalimumab improves survival, functional engraftment and A9 enrichment of purified engrafted postmitotic dopamine neurons

Next, we assessed whether a widely used, FDA-approved TNFa neutralizing antibody, adalimumab (Humira^®^), can enhance the survival of CD49-low/CD184-high purified, postmitotic dopamine neurons. Adalimumab is known to bind soluble and transmembrane bound TNFa (McCoy and Tansey, 2008) and is approved for the treatment of autoimmune disorders such as rheumatoid arthritis and plaque psoriasis. We found that a single co-injection of adalimumab, added to the cell suspension at grafting significantly improved the survival of CD marker purified dopamine neurons (**Figure 7A and Figure S7C**). Stereological counts of NURR1::GFP+ cells at 1 month post grafting found 12,423 (mean) ± 1,859 (S.E.M) in adalimumab treated grafts per 100,000 cells injected versus 6,057 (mean) ± 378 (S.E.M) in PBS treated neurons; p = 0.01, paired t-test, n = 5)] (**Figure 7B**). The volume of the adalimumab treated graft was 0.1175 mm^3^ (mean) ± 0.01 (S.E.M) mm^3^ versus 0.05325 mm^3^ (mean) ± 0.003 mm^3^ (S.E.M) mm^3^ per 100,000 cells in PBS-treated grafts (p = 0.0085, paired t-test, n = 5).

**Figure 7.**
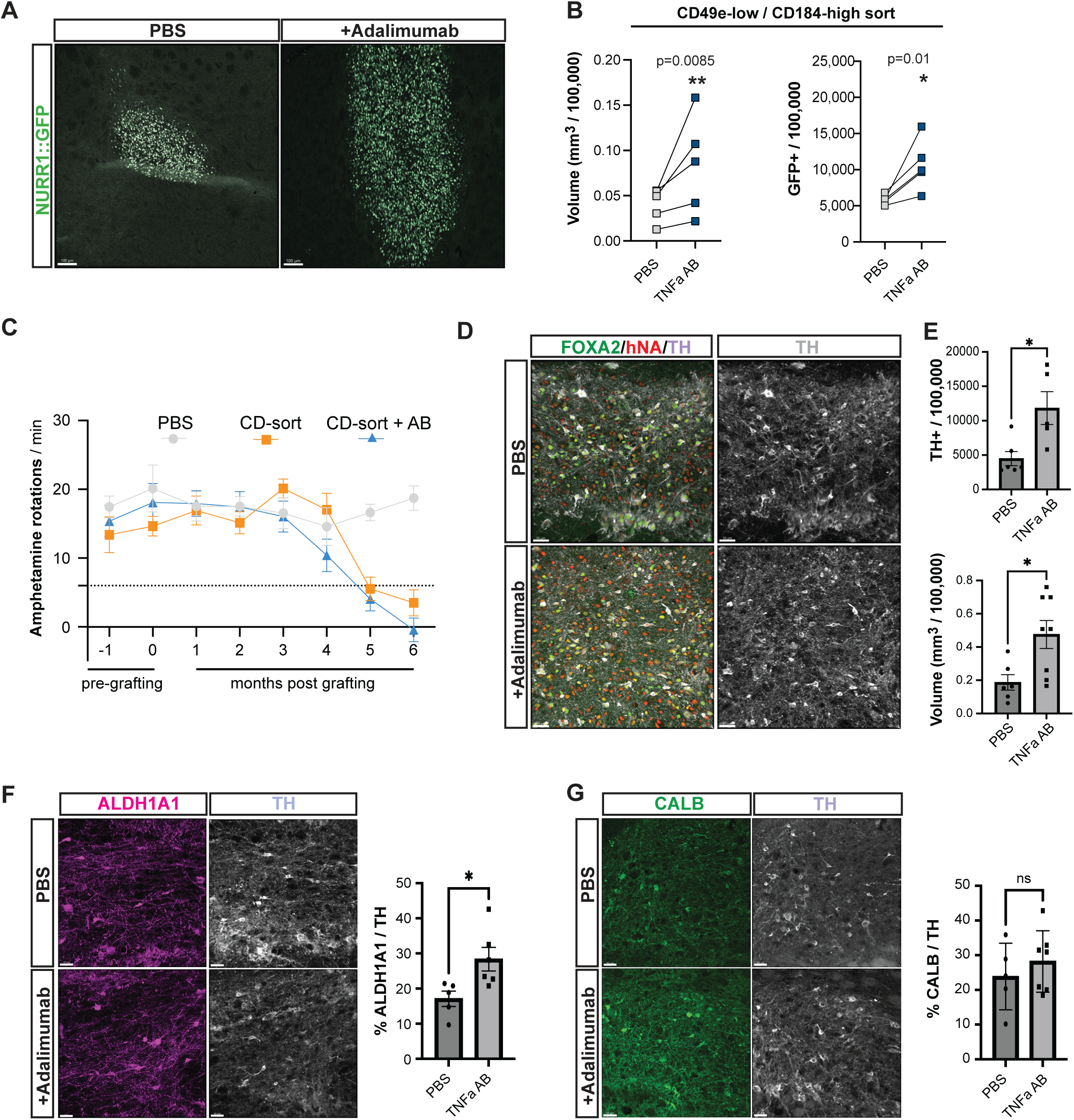
Clinically relevant TNFa neutralizing antibodies functionally improve the survival of postmitotic dopamine neuron following implantation. (**A**) Representative immunofluorescence image of CD49e-low/CD184-high double sorted dopamine neurons either co-injected with PBS or TNFa blocking antibody, adalimumab. Scale bar = 100 μm. (**B**) Stereological cell counts and volume quantification of the surviving dopamine neurons at 1 month post transplantation using NURR1::GFP, n = 5. **p<0.01, *p<0.05 (paired t-test) (**C**) D-amphetamine induced rotation assay in grafted PD mouse model carrying unilateral 6-OHDA lesion. The three injection groups are: PBS, CD sorted neurons (CD-sort), and CD sorted neurons co-injected with adalimumab (CD-sort +AB). (**D**) Representative immunofluorescence images of human grafts that are highly enriched with floor-plated derived dopamine neurons marked by hNA, FOXA2, and TH for each group, scale bar = 50 μm. (**E**) Stereological analysis of the number (using optical fractionator) and volume (cavalier estimator) of the surviving dopamine neurons based on TH expression at 6 months post transplantation. *p<0.05 (paired t-test). (**F, G**) Representative immunofluorescence image and quantification of portion of ALDH1A1 demarking A9 subtype (**F**) and CALB1 demarking A10 subtype (G) dopamine neurons population (TH+) in 6 months old graft.

Finally, we sought to examine whether CD sorted postmitotic dopamine neurons, with or without adalimumab treatment, are functional in rescuing amphetamine-induced rotation behaviors in a 6-OHDA treated mice, which represents a widely used preclinical model for treating PD motor symptoms. We observed a robust functional recovery, regardless of adalimumab exposure, at 6 months post grafting. However, adalimumab-co-injected dopamine neurons showed a trend towards an earlier and more complete functional rescue (**Figure 7C**).

Histological analyses showed that both groups give rise to highly enriched neuronal population *in vivo* at 6 months post implantation (**Figure S7A and B**). However, TNFa inhibition via adalimumab resulted in significantly increased total numbers of surviving dopamine neurons as well as overall graft size (**Figure 7D and E**). Compared to the 1-month survival data (**Figure 7B**), we postulate that most dopamine neurons at 1 month survive at 6 months with > 80% giving rise to TH positive (+) neurons. This is consistent with our previous report, demonstrating nearly 100% conversion of NURR1::H2B-GFP populations into TH expressing neurons *in vitro* based on a single cell RT-qPCR assay (Riessland et al., 2019). Similar to the results with p53 KO cells, we observed that adalimumab treatment results in significant increase in the proportion of ALDH1A1+ A9 dopamine neuron subtype without affecting the fraction of CALB1 expressing A10 dopamine neuron subtype (**Figures 7F and G**).

These results offer an important proof of concept by addressing two major, unresolved challenges in the field, which is the ability to routinely purify postmitotic dopamine neurons suitable for *in vivo* transplantation without the need for genetic modification, and the identification of key regulators of dopamine neuron death post grafting. The combination of those two strategies results in a clinically relevant solution to overcome limited dopamine neuron graft survival using an FDA approved drug and enables the functional engraftment of purified postmitotic dopamine neurons.

## Discussion

The main gene identified in our *in vivo* CRISPR/Cas9 screen for dopamine neuron graft survival was *p53*. p53 is a master regulator of diverse cellular processes ranging from tumor suppression to serving as a guardian of cell fate identity and reprogramming to sensing cellular stresses related to DNA damage, oxidative stress, or ischemic injury (Hafner et al., 2019). p53 has been recently implicated as a candidate factor for driving the vulnerability of human substantia nigra dopamine neurons during PD pathogenesis based on single cell data from human patients (Kamath et al., 2021; Kamath et al., 2022; Monzon-Sandoval et al., 2020). Furthermore, p53 may serve as a more general signaling hub of neuronal cell death across neurodegenerative disorders (Maor-Nof et al., 2021). Our initial screen identified several hits in addition to p53 as limiting *in vivo* dopamine neuron survival. Several of those hits can be linked to the TNFa/p53/apoptotic pathway described here including *TNFRSF11B, BBC3, BCL2L11, CASP2, CASP9.* Other hits such as *SLC7A11, IL18* or *TLR4* are associated with ferroptosis or neuroinflammatory responses. Future studies will be required to further validate these additional hits and to explore whether they act independently or in concert with the TNFa/p53/apoptotic axis.

Our study focused on cell intrinsic factors driving dopamine neuron death by screening for genes limiting survival directly in purified dopamine neurons without directly manipulating host-related responses. Similarly, our gene expression data focuses on changes in grafted and re-isolated dopamine neurons *via* either bulk or single cell-based RNAseq analyses. On the other hand, our histological data present a cascade of host cell-related cellular responses at the graft site including the recruitment of neutrophils, microglial and vascular cells that enter the graft core by 3 dpt, and the activation of inflammatory programs in both astrocytes and microglia. Temporal data suggest that those host responses occur in response to rather than mediating dopamine neuron death. However, our work does not rule out a more direct role of the host response on graft survival and function, as glial cell activation has been proposed as key driver of neurodegeneration including the loss of dopamine neuron in the substantia nigra of human PD patients (Liddelow et al., 2017). While grafting-related dopamine neuron death in our study subsided completely by 7 dpt, immune activation is known to persist in dopamine neuron grafts for months if not years post-surgery (Winkler et al., 2005). It will be interesting to explore whether blocking dopamine neuron death *via* p53 or TNFa inhibition also impacts the extent of such long-term, graft-related neuroinflammatory responses, though in the postmitotic dopamine neurons grafts presented here, there is little evidence of a persistent host inflammatory response at 6 months post grafting.

Gene expression data suggest that the grafted dopamine neurons are a source of TNFa following transplantation, possibly in response to injury-related damage. TNFa is known to be secreted in response to hemorrhagic, ischemic or traumatic injury in the brain (Tuttolomondo et al., 2014), events commonly associated with cell transplantation. However, host glial cells could also contribute to TNFa mediated activation of NFκB and p53 in dopamine neurons. To address this point, future studies could involve grafting into TNFa KO mice to directly assess the contribution of host-derived TNFa. In either case, it will be important to determine the upstream events leading to the specific induction of TNFa *in vivo,* as we did not observe a similar TNFa/NFκB/p53 cascade upon replating the identical dopamine neuron population *in vitro*.

While p53 loss or treatment with TNFa inhibitory antibody led to a dramatic, 3-10 fold increase in dopamine neurons survival (from 2-5% to 15-25%), we were unable to rescue all the grafted dopamine neurons. We currently do not know whether the remaining dopamine neuron death is caused by incomplete suppression of p53-dependent or the action of p53-independent mechanisms. At 1dpt, we report the induction of p53 in 40% of the grafted neurons but the presence of TUNEL positive apoptotic dopamine neurons representing 60% of the cells. Therefore, both p53 dependent and independent mechanisms may contribute to early dopamine neuron death. Transcriptomic analysis implicates hypoxia in grafted neurons as a potential additional contributing factor. Interestingly, in a 3D organoid model that recapitulates hypoxic brain injury, cell death was independent on the activation of p53/CC3 activation (Pasca et al., 2019) suggesting that hypoxia could serve as an independent driver of dopamine neuron death.

The use of a genetic reporter line to purify postmitotic dopamine neurons allowed us to perform a survival screen that avoids confounding factors related to cell proliferation or non-autonomous signaling among grafted cells. In addition, this approach sets the stage for grafting postmitotic dopamine neurons for translational applications, based on our previous work identifying the NURR1/NR4A2 stage as particularly suitable to achieve dopamine neuron survival *in vivo* (Ganat et al., 2012). The CD surface marker-based selection strategy presented here enables dopamine neuron enrichment at the NURR1 stage without the need for establishing genetic reporter lines. Therefore, this strategy should be compatible for future translational applications in dopamine neuron replacement therapy in PD. Furthermore, the same strategy can facilitate the use of purified dopamine neuron preparations for *in vitro* studies such as human iPSC-based disease modeling efforts given the well-known challenges for reliably generating specific neuron subtypes across many cell lines and laboratories (Schwartzentruber et al., 2018; Volpato et al., 2018).

In conclusion, our data suggests a novel approach for cell therapy that may enable the transplantation of postmitotic dopamine neurons with improved survival to maximize safety in PD patients by avoiding the introduction of off-target cells and avoiding the need for grafting dopamine neuron precursor cells that retain some proliferative capacity *in vivo* and may yield various proportions of undesired graft-derived glial lineages in long-term grafts. It will be exciting to explore whether the same approach can be adapted for other ongoing cell therapy efforts within or outside the CNS, as the limited *in vivo* survival is a major impediment across many hPSC-based cell therapy paradigms.

## Limitation of the study

Our study is based on CRISPR/Cas9 screening for a limited set of candidate genes related to mechanisms of cells death. Therefore, we cannot rule out that other genes, not included in our screen, may have similar or stronger effects than p53 reported here. Using our experimental setup, it was not feasible to perform a whole-genome screen, as this would require the injection of ∼ 1 billion dopamine neurons across ∼1,000 mouse brains. Furthermore, our study specifically addresses intrinsic regulators of dopamine neuron death. Additional strategies that could be explored to address extrinsic factors include the transient elimination of host microglia using PLX3379 treatment, a strategy used to study the impact of microglia on amyloid beta pathology (Spangenberg et al., 2016) or blocking of the host neutrophil response using monoclonal antibodies (Nguyen et al., 2017). Finally, despite the larger number of dopamine neurons surviving in adalimumab treated grafts, we observed functional recovery in both adalimumab and PBS groups. This is likely due to a saturation effect with limited threshold of surviving dopamine neurons required to normalize amphetamine-induced rotation behavior. Detailed future studies will be required to perform dose-response studies across several pre-clinical PD models to demonstrate the full potential of this approach for eventual human translation.

## Acknowledgements

We thank all the Studer Lab members for insightful comments and feedback on this project. We further thank Stefan Iron and Rui Gardner for setting up the cell surface screening. We would also like to thank the Flow Cytometry core, the Molecular Cytology core, the Integrated Genomic Operation at MSKCC for outstanding technical support. We acknowledge the use of the Integrated Genomics Operation Core, funded by the NCI Cancer Center Support Grant (CCSG, P30 CA08748), Cycle for Survival, and the Marie-Josée and Henry R. Kravis Center for Molecular Oncology. This work was supported in part through grants 1R01 NS118067-01A1 (L.S., D.B.), 1R01 NS124735-01A1 (M.R.), NIH T32 GM132038 (F. C), and support from the JPB foundation to L.S. The research was funded, in part, by Aligning Science Across Parkinson’s [Grant number: ASAP-000472] through the Michael J. Fox Foundation for Parkinson’s Research (MJFF). T.W.K was supported in part by a Druckenmiller fellowship from the New York Stem Cell Foundation.

## Author Contribution

L.S., T.W.K., and S.Y.K., conceived and designed the experiments, performed data analysis and interpretation, and wrote manuscript.

T.W.K., S.Y.K., M.R., B.K., performed dopamine neuron differentiation, *in vivo* transplantation, CRISPR screening, histological studies, cell surface marker screening.

N.S., provided a construct for CRISPR screen.

H.W.C., F.C., performed bioinformatics analyses and interpretation of data. D.B., supervision of computational analysis and interpretation of data.

M.V.R., H.W.C., S. M., and R. G., performed CRISPR/Cas9 screening analyses. All authors provided feedback in editing the manuscript.

## Figure legends to Supplementary Figures

**Figure S1. Derivation and validation of NURR1:GFP sorted dopamine neurons for CRISPR/Cas9 screening *in vitro* and *in vivo***.

(**A**) Immunofluorescent staining of dopamine neuron markers, NURR1:GFP, FOXA2, and TH, in NURR1:GFP sorted cells two weeks post sorting (day 40). Scale bar = 100 μm. **(B)** Immunofluorescent staining of dopamine neuron markers, TH and NURR1::GFP in the graft 1 month post transplantation of day 25 NURR1:GFP sorted dopamine neurons. Scale bar = 100 μm. **(C)** Transduction efficacy and isolate transduced hPSC containing pooled lentiviral sgRNAs expressing Tomato with MOI = 0.35, which indicates single copy sgRNA integration per cell. **(D)** Immunofluorescent staining of TH, NURR1::GFP, and gRNA::Tomato at day 40 dopamine neurons post sorting with GFP and Tomato at day25 with/without dox treatment from day16 to day25. Scale bar = 50 μm. **(E)** Ablation of the Tomato signal in hPSC-derived postmitotic dopamine neurons at day25 after dox exposure from day16 to day25 (dox 1ug/ml). Scale bar = 200 μm. **(F)** Fluorescence expressions in graft 1 month post transplantation (upper). PCR analysis for detecting a human PTGER2 gene from genomic DNA, isolated from dissected tissue around graft region (lower). Scale bar = 500 μm. **(G, H)** Overall representation of sgRNA across all condition by next-generation sequencing (NGS) from genomic DNA of *in vitro* cultured cells (day 16 and day25) and an *in vivo* grafted cell from library **(G)** and more restricted library#2 **(H).**

**Figure S2. Increased survival of P53 KO dopamine neurons in graft exhibit dopamine neuron identity and needle injection of dopamine neuron does not induce P53 and P53 downstream genes**.

(**A**) Immunofluorescent staining of NURR1:GFP, sgP53RNA-Tomato, and FOXA2 and TH at day40 dopamine neuron post sorting with GFP and Tomato at day25 with/without dox exposure from day16 to day25. Scale bar = 100 μm. (**B-E**) Representative immunofluorescence image for dopamine markers, such as FOXA2, TH, and NURR1-GFP (**B-D**) as well as a proliferation marker, hKI67 (**E**) expression in surviving p53 WT and KO dopamine neurons in the graft (WT;-DOX, KO; +DOX). (**F**) qRT-qPCR analysis of P53 and P53 downstream genes (p21 and PUMA) before and 1 hour after needle injection of NURR1:GFP sorted dopamine neuron.

**Figure S3. Time-course analysis of host neuroimmune cells after transplantation near the graft site**.

**(A)** Immunofluorescence analysis of IBA1 (upper) and GFAP (lower) following transplantation (4hrs, 24hrs, 72hrs, 7days). Scale bar = 50 μm. **(B)** H&E staining of the graft. Scale bar = 100 μm. **(C)** Immunofluorescence and immunohistochemistry analysis for Ly6G to examine neutrophils at the graft site. Left panels are examined at 12-hour post transplantation (hpt) and yellow arrows indicate Ly6G positive cells nearby the grafted cells positive for FOXA2. Right panels are examined at 72hpt. Ly6G positive cells marked by dark brown staining infiltrate within the graft. Scale bar = 50 μm. **(D)** Immunofluorescence analysis of fiber outgrowth pattern from the graft after implantation using STEM 121. Fiber extension begins at 24 hpt. Scale bar = 50 μm.

**Figure S4. Analysis of bulk RNA-seq from sorted cells *vs.* 1 day cultured cells post sorting *vs.* 1 day grafted cells post sorting, and characterization of 1 day cultured cells post sorting**.

(**A**) Dendrogram of the cells among sorted (Day 0), *in vitro* cultured (Day 1 Culture), and *in vivo* grafted dopamine neurons (Day 1 Graft) from bulk RNA-seq, demonstrating agreement among the replicate samples and distinct signature of the day 1 grafted samples. (**B**) Pathway enrichment analysis identified mTORC1 signaling as upregulated categories in day 1 cultured than day 1 grafted samples. (**C**) Heatmap analysis from total RNA-seq for enriched TNFa-NFκB related genes in the grafted dopamine neurons versus the sorted and cultured dopamine neurons. (**D**) Immunofluorescence staining of NURR1::GFP, P53, and NFkB-p65 at day 26 dopamine neuron in culture 1 day post sorting with GFP shows no nuclear induction of NFkB and p53. Scale bar = 100 μm.

**Figure S5. scRNA-seq analysis from p53 WT and KO dopamine neuron grafts 1 day post implantation**.

**(A)** Clustering distribution of p53 WT and p53 KO of grafted dopamine neurons 1 day post transplantation from scRNA-seq. **(B)** Histograms of fraction of cells expressing MAP2, MKI67, and TH positive cells in p53 WT and KO. **(C)** UMAP plots of MAP2, PBX1, and MKI67 in p53 WT and p53 KO neurons from scRNA-seq. **(D)** Volcano plot of differentially expressed genes in p53 WT versus p53 KO from the clusters of 1, 2, and 4.

**Figure S6. Characterization of cell surface (CD) marker sorted cells enriching NURR1::GFP dopamine neuron *in vitro* and *in vivo***.

**(A)** FACS analysis of NURR1::GFP neuron population with indicated CD markers, demonstrating 3 CD markers (49e, 99, and 340) are negatively whereas 2 CD markers (171 and 184) are positively enriched to NURR1::GFP populations. **(B)** Immunofluorescence analysis of grafted cells from double CD49-low/CD184-high sorted and unsorted cells with FOXA2 and a human proliferation marker (hKI67) at 1 month post transplantation (left) and quantification of KI67 positive cells within the grafts (right).

**Figure S7. Innervation of grafts from CD marker sorted dopamine neurons co-injected either PBS or TNFa inhibitor, adalimumab, at 6 months**.

(**A, B**) Representative image of extensive innervation of CD marker sorted dopamine neuron following transplantation co-injected with PBS or adalimumab towards the host striatum as indicated by TH (A) and hNCAM immunofluorescent staining (**B**). Scale bar = 500 μm. (**C**) Immunofluorescence assay to examine the spread of adalimumab within the brain. Anti-human IgG1 Alexa fluorophore 555 was probed to detect the presence of adalimumab comparing adalimumab co-injected neurons *vs.* PBS injected neurons at 24 hpt. Red signal indicates a specific detection of the human monoclonal antibodies only present in adalimumab injected group.

## METHOD

### Cell line

H9 (WA-09, passage 40-60) human pluripotent stem cell (hPSC) line was employed throughout the study, which engineer to generate *Nurr1::GFP* reporter hPSC and doxycycline-inducible CRISPR/Cas9 expression in the *Nurr1::GFP* hPSC line (iCas9/*NURR1::GFP* hPSC) as well as iCas9/*NURR1::GFP* hPSC lines containing sgRNA-pool libraries and sgRNA for dTomato and p53. hPSCs were grown in feed-free conditions on vitronectin (VTN-N; Thermo Fisher Scientific)-coated dishes in E8-essential medium and maintained at 37℃, 5% CO_2_. Tri-I (MSKCC, Weil-Cornell, Rockefeller University) Embryonic Stem Cell Oversight (ESCRO) approved this study.

*Construction of Nurr1::GFP and inducible expression of CRISPR/Cas9 in Nurr1:GFP hESC lines.* Generation of Nurr1::GFP hESC line was previously described (Riessland et al., 2019). Briefly, stop codon of endogenous NR4A2 (Nurr1) was replaced by EGFP expression cassette (P2A-H2B-PgkPuro) by using a CRISPR/CAS9-mediated knock-in approach. The resulting *NURR1:GFP*^+^ cells almost express TH (a mature mDA marker; 98%) based on single cell qRT-PCR^9^. To generate doxycycline-inducible CRISPR/Cas9 expression in the *Nurr1::GFP* hPSC line (iCas9/*NURR1::GFP* hPSC line), a pair of TALEN, Neo-M2rtTA donor, and Hygro-Cas9 donor (Addgene#86883) (Gonzalez et al., 2014), targeting to an *AAVS* locus, were transfected into the Nurr1::GFP hPSC using Nucleofector (Lonza, B-016 program) in AMAXA machine and stable cell line was generated following a published protocol (Gonzalez et al., 2014). Briefly, 2 days after transfection, neomycin and hygromycin (100ug/ml) were treated for 1 week and picked clonal expanded hPSC. Inducible expression of Cas9 in each clone was confirmed by immunofluorescent staining with a Cas9 antibody after Dox (1ug/ml) exposure for 3 days.

### Single-strand guide RNA (sgRNA) Design and Cloning

sgRNA sequences for pool library were identified by Guidescan (MSKCC) and sgRNA oligos were synthesized on-Chip (Agilent). Synthesized oligos were PCR amplified and amplicons were restriction cloned into SGL40C.EFS.dTomato (<u>Addgene#89395</u>). Library representation was assessed by NGS (Illumina). Individual sgRNA for dTomato and p53, designed by web-based tool (http://crispor.tefor.net) and using Guidescan (Perez et al., 2017)(MSKCC) subsequently, were restriction cloned into SGL40C.EFS.dTomato plasmid vector (<u>Addgene#89395</u>).

### Lentiviral production and transduction

SGL40C.EFS.dTomato vector containing sgRNA for libraries, dTomato, and p53 was co-transfected with packing vectors, psPAX2 (Addgene#12260) and pMD2.G (Addgene#12259) into HEK293T cell using Xtream Gene 9 transfection reagent (Sigma). The virus was collected after 2 days of transfection, and infected into the iCas9/*NURR1::GFP* hPSC. 2 days post infection, dTomato expressed hPSCs were sorted using flow cytometry associated cell sorting (FACS) in Flow Cytometry Core Facility at MSKCC. The sorted hPSCs were cultured and maintained for subsequent experiments until use.

### Dopamine neuron differentiation from hPSC

Human PSCs were directed differentiated into midbrain dopamine neuron with a clinically relevant protocol from a previously reported study (Kim et al., 2021). At day 25 of differentiation from hPSCs iCas9/*NURR1::GFP* hPSC lines containing sgRNA-pool libraries and sgRNA for dTomato and p53 hPSCs, GFP and dTomato expressed dopamine neurons were sorted using flow cytometry associated cell sorting (FACS) in Flow Cytometry Core Facility at MSKCC. The sorted hPSCs were either injected into mice or cultured for subsequent experiments until use. Furthermore, double sorting approach with CD49e-low and CD184-high was applied to enrich post-mitotic dopamine neuron at day 25 differentiation derived from hPSC using FACS, and sorted cells were used for transplantation and *in vitro* culture. TNF-a neutralizing antibody was employed either to co-injection (1mg/ml) or *in vitro* cultured dopamine neuron (10ug/ml). 100ng/ml of TNFa was added into NURR1::GFP sorted dopamine neurons to model TNFa-NFκB - p53 axis in culture.

### sgRNA barcode Sequencing and Analysis to identify targets

Cell Pellets at each desired time point were lysed, and genomic DNA was extracted (Qiagen) and quantified by Qubit (Thermo Scientific). A quantity of gDNA covering 1000X representation of sgRNAs was PCR amplified to add Illumina adapters and multiplexing barcodes. Amplicons were quantified by Qubit and Bioanalyzer (Agilent) and sequenced on Illumina HiSeq 2500. Sequencing reads were aligned to the screened library and counts were obtained for each gRNA at MSKCC Gene Editing and Screening Core Facility. The resulting single end reads were checked for quality (FastQC v0.11.5) and processed using the Digital Expression Explorer 2 (DEE2) workflow (Chen et al., 2015). Adapter trimming was performed with Skewer (v0.2.2) (Wang et al., 2021a). Further quality control done with Minion, part of the Kraken package (Michels et al., 2020). Differential gRNA hits were identified using EdgeR, a Bioconductor package (Robinson et al., 2010), to identify the primary hits. We used the Trimmed Mean of M-values for normalization and the glmQLFTest / F test for statistic tests. Additionally, we used the camera analysis function from EdgeR for gene-level analysis as previously described (McCarthy et al., 2012; Robinson et al., 2010). To calculate the correlation between the screen samples, quantifications were normalized by the estimateSizeFactorsForMatrix of DESeq2 (Love et al., 2014) using only the non-targeting control and safe harbor probes. Pearson correlation of the pairwise comparison was plotted using an R package pheatmap (https://CRAN.R-project.org/package=pheatmap) (Kolde, 2018).

### Cell preparation for survival surgery and intracranial transplantation

hESCs-derived mDA neurons that were sorted based on either NURR1::H2B-GFP or CD49e low and CD184 high were resuspended in 100,000 ± 10,000 cells/µL in neurobasal medium with 200 mM L-glutamine and 100 mM ascorbic acid (AA) transplantation medium (without human albumin or kedbumin 25%) (Kim et al., 2021). Unless specified, 3-4 µL of sorted neurons were injected at the rate of 0.5 – 1 µL per minute (1 µL per deposit) into the striatum of wild-type (unlesioned) 6 to 8-week-old male NSG mice ([AP] +0.5mm, [ML] +/-1.8mm, [DV] −3.4 to −3.3 mm from dura). Each surgery did not exceed more than 30 minutes per animal and the entire surgery time was within 8-10 hours post cell preparation. For the short-term 1-month survival study, p53 WT (-dox) *vs.* KO (+dox) NURR1::GFP+ dopamine neurons or CD + PBS vs. CD + adalimumab was bilaterally engrafted into the striatum of the same mouse brain to reduce variability between animals. For other studies including time course or behavior studies, cells are engrafted unilaterally. For the time course experiments, the mice were euthanized and used for immunohistochemistry analysis at designated time points (4hrs, 1day, 3days, and 7days post engraftment). For *in vivo* CRISPR screen, maximum loadable amount of the cells is implanted with cell density of 200,000 ± 10,000 cells/µL (4 µL total in each striatum) in order to have enough representations of the guide RNAs. All animal work was performed under the approval of Institutional Animal Care and Use Committee (IACUC) at Memorial Sloan Kettering Cancer Center.

### DNA extraction from the xenograft sample and detection of human DNA

Grafted dopamine neurons at 1 month were isolated from thick tissue slices based on NURR1::GFP and sgRNA::Tomato expressions and extracted genomic DNAs using the DNeasy Tissue kit according to the manufactural protocol (Qiagen #69556). To detect human DNA from the extracted genomic DNA, PCRs were performed to detect human and mouse cells with human and mouse specific primers for *PTGFR2* and *ptgfr2* subsequently using the Q5 High-Fidelity PCR kit (NEB #M0493S). Primer sequences are below.

Human PTGFR2 primer 5’: gctgcttctcattgtctcgg

Human PTGFR2 primer 3’: gccaggagaatgaggtggtc

Mouse ptgfr2 primer 5’: cctgctgcttatcgtggctg

Mouse ptgfr2 primer 3’: gccaggagaatgaggtggtc

### 6-OHDA mouse model

Briefly, adult female and male NSG mice (6-12 weeks) were anesthetized with 1%–2% isoflurane mixed in oxygen. 1 µL 6-OHDA (3 mg/ml, in saline with 1% ascorbic acid) was directly injected into the right side of substantial nigra (anterior-posterior [AP] = − 2.9 mm, lateral [ML] = 1.1 mm, vertical [DV] = 4.5 mm, from dura) with rate of 0.5 – 1 µL per minute to generate unilateral toxin Parkinsonian mouse model. Animals with amphetamine-induced rotation at > 6 rotations per min were selected for cell transplantation 4 weeks after 6-OHDA-lesion surgery. Animals were randomly grouped and transplanted with CD sorted neurons vs. CD sorted neurons + adalimumab vs. PBS (sham surgery control).

### d-amphetamine induced rotation behavior assays

Amphetamine-induced rotation tests were performed twice before transplantation, and 1, 2, 3, 4, 5, 6 months after transplantation. The mice were injected intraperitoneally with a d-amphetamine in saline (Sigma, 10 mg/kg). After 10 minutes, the rotation behavior was recorded for 30 minutes, and the total rotations were automatically counted by Ethovision XT 11.5. The data were presented as ipsilateral – contralateral rotations per minute.

### Cell and Tissue Processing for immunofluorescence (IF) labeling

Histology on tissues from mice was performed on frozen sections from xenografts. Mice were anesthetized with pentobarbital and transcardically perfused using heparinized (10U/mL) PBS (pH 7.4), followed by 4% paraformaldehyde in PBS. The liquid is administered using peristaltic pump to control the rate of the solution delivery to the system. Tissues were post-fixed in ice cold 4% paraformaldehyde for 18 hours sharp and transferred to 30% sucrose until the tissue sink (typically 3-6 days post-), followed by snap freezing in O.C.T (Fisher Scientific, Pittsburgh, PA) or Neg-50 (Thermo Scientific). Brain tissues are all sectioned in 30 μm thick coronal sections using a cryostat and mounted onto a Superfrost plus microscope slides (Fisher Scientific). All the slides are stored at −80℃ for long-term until use. To process for immunolabeling, tissues are washed twice with 1x PBS, followed by permeabilization in in 0.5% Triton X-100 in PBS for 10 minutes. Living cells in culture were directly fixed in 4% paraformaldehyde for 18 min, followed with 10 min permeabilization in 0.5% Triton X-100 in PBS. For labeling, cells or tissue sections were immuno-stained with primary antibodies of interest in 2% BSA in 0.25% Triton X-100 in PBS at 4°C overnight. Next day appropriate Alexa Fluor secondary antibodies are conjugated at room temperature for 1hr at a dilution of 1:1,000. The information for primary antibodies and secondary antibodies is provided in **supplemental Table 4**. Nuclei were counterstained by DAPI.

### Tissue immunohistochemistry (IHC), TUNEL, and H&E stain

H&E and IHC on tissues from mice was performed on FFPE (formalin fixed paraffin embedded) sections from xenografts. Mice were anesthetized with pentobarbital and transcardically perfused using heparinized (10U/mL) PBS (pH 7.4), followed by 10% formalin. The liquid is administered using peristaltic pump to control the rate of the solution delivery to the system. Tissues were post-fixed in 10% formalin 24-48 hours at room temperature (can stay up to a week in 10% formalin) and directly transferred to 70% ethanol. Histology was performed by HistoWiz Inc. (<u>histowiz.com</u>) using a Standard Operating Procedure and fully automated workflow. Samples were embedded in paraffin, and sectioned at 4μm. Immunohistochemistry was performed on a Bond Rx autostainer (Leica Biosystems) with heat-mediated antigen retrieval using Epitope Retrieval Solution 1 (Leica Biosystems). Primary antibodies used were rabbit polyclonal total NFKB (Cell Signaling, CST8242, 1:300), Cleaved Caspase-3 (Cell signaling, CST9661, 1:300), p53 (CM5P, 1:500), and p-NFkB (GTX55113, 1:5000) followed by anti-rabbit HRP conjugated polymer system. Bond Polymer Refine Detection (Leica Biosystems) was used according to the manufacturer’s protocol. After staining, sections were dehydrated and film coverslipped using a TissueTek-Prisma and Coverslipper (Sakura). Whole slide scanning (40x) was performed on an Aperio AT2 (Leica Biosystems).

For TUNEL: Standardized conditions using the Promega DeadEnd Colorimetric Detection System (G3250), Enzyme Digestion for 10 minutes, using the Leica Bond Polymer Refine Detection Kit (DS9800). For H&E, staining was performed on Sakura Autostainer. Briefly, deparaffinize the slides in 2 changes of xylene, 2 changes of 100% alcohol, 1 change in 95% alcohol, then wash with water. Place slides in this sequence: hematoxylin, a rinse with water, define solution, a rinse with water, bluing agent solution, rinse with water, 95% alcohol, eosin, and 95% alcohol. Finish with two changes of 100% alcohol and two with xylene.

### Stereological analysis

For short-term survival studies, unbiased stereological counts of *NURR1::H2B-GFP* DA neurons within the striatum (AP +0.5, ML +/- 1.8, DV −3.4 to −3.3 from dura) were performed using stereological principles and analyzed with StereoInvestigator software (Microbrightfield, Williston, VT, USA), as previously described (Piao et al.,). For long-term behavioral studies, unbiased stereological counts of TH positive DA neurons within the striatum were performed. The tissue was embedded in O.C.T. or Neg-50 and sections are sliced at 30 µm. Optical fractionator sampling (Boyce et al., 2010) was carried out with an Olympus BX61 microscope equipped with a motorized stage and Olympus 40x objective lens objective. Graft region was outlined on the basis of NURR1::GFP immunolabeling, with reference to a coronal atlas of the mouse brain (Franklin and Paxinos 1997). Every 3^rd^ - 10^th^ section (depending on the total thickness of the graft) from the beginning of the graft to the end of the graft was randomly and systematically selected for analysis. For each tissue section analyzed, section thickness was assessed in each sampling site and guard zones of 1 μm were used at the top and bottom of each section. Pilot studies were used to determine suitable counting frame and sampling grid dimensions prior to counting to achieve enough statistical power and low Gunderson coefficient variance.

The following stereological parameters were used in the final study: for optical fractionator probe, 65 μm x 65 μm optical dissector, 100 μm x 100 μm (or 10% of ROI) SRS, 20 μm optical dissector height and 1 μm guard zone; for cavalier estimator probe, 50 μm x 50 μm grid spacing, 0-degree grid rotation, and 30 μm section cut thickness. For analysis, at least 2-8 sections were evaluated for analysis. Gundersen coefficients of error for all conditions were less than 0.1. Stereological estimations were performed with the same parameters for all experimental conditions, p53 KO vs. WT (NURR1::GFP sort) or TNFa monoclonal antibodies treatment vs. PBS (CD sort).

### RNA extraction and Real time quantitative reverse transcription-PCR (qRT-PCR)

Total RNA samples were prepared from the cells and DNase I treated using TRIzol according to the manufacturer’s instructions. Delta-delta-cycle threshold (ΔΔCT) was determined relative to GAPDH levels and normalized to control samples. Error bars indicate the standard deviation of the mean from three biological replicates. The sequences of qRT-PCR primers are shown below.

Human GAPDH primer 5’: ATGTTCGTCATGGGTGTGAA

Human GAPDH primer 3’: AGGGGTGCTAAGCAGTTGGT

Human FOXA2 primer 5’: CCGACTGGAGCAGCTACTATG

Human FOXA2 primer 3’: TACGTGTTCATGCCGTTCAT

Human NURR1 primer 5’: CGCTTCTCAGAGCTACAGTTAC

Human NURR1 primer 3’: TGGTGAGGTCCATGCTAAAC

Human PUMA primer 5’: CCTGGAGGGTCCTGTACAATCT

Human PUMA primer 3’: TCTGTGGCCCCTGGGTAAG

Human P53 primer 5’: GTACCACCATCCACTACAACTAC

Human P53 primer 3’: CACAAACACGCACCTCAAAG

### High throughput cell surface marker screen and enrichment of dopamine neuron with using cell surface markers

Dopamine neuron differentiated cells at day25 from the NURR1::GFP reporter hESC were single cell suspended in flow cytometer staining buffer (PBS containing 2% bovine serum albumin). The cells were stained with 387 cell surface (CD) markers (0.2M cells per a CD marker) for 30 min on ice in the dark. After 3 times washing with PBS, cells were co-stained with DAPI. All staining for the screen was done in 96 well plates. Data collection using a flow cytometer to identify CD markers to enrich GFP positive population was performed by the MSKCC Flow Cytometry core facility. For enrichment of dopamine neuron with using CD markers, 49e and 184, day25 cells were stained with 49e and/or 184, followed by isolation of 49e weak, 49e weak and 174 high, and 49e weak and 184 high expressed cells via FACS at the MSKCC Flow Cytometry core facility.

### Protein isolation and western-blot analysis

Cells treated with PBS or monoclonal antibodies against TNFa were harvested and lysed in the following lysis buffer (RIPA buffer, 1:1000 Halt^TM^ Protease and Phosphatase Inhibitor cocktail (Thermo Fisher Scientific). After cells are resuspended in a lysis buffer, cells are sonicated for 3×30 seconds at 4°C. Supernatant was collected upon 15 minutes centrifugation >15,000 rpm at 4°C and protein concentration was measured using Precision Red Advanced Protein Assay (Cytoskeleton). Equal amounts of protein (20 micrograms) were boiled in NuPAGE LSD sample buffer (Invitrogen) at 95°C for 5 minutes and separated using NuPAGE 4%-12% Bis-Tris Protein Gel (Invitrogen) in NuPAGE MES SDS Running Buffer (Invitrogen). Proteins were electrophoretically transferred to a nitrocellulose membrane (Thermo Fisher Scientific) with NuPAGE Transfer Buffer (Invitrogen). Blots were blocked for 60 minutes at RT in 5% nonfat milk (Cell Signaling) in TBS-T + and incubated with respective primary antibody at 4°C overnight. The following primary antibodies were used: mouse-anti-GAPDH (6C5) (Santa Cruz, 1:1,000); mouse anti-p53 (DO-1) (Santa Cruz, 1:1,000); rabbit anti-tyrosine hydroxylase (Pel Freeze, 1:1000). Primary antibodies were detected using the secondary anti-rabbit IgG HRP-linked (Cell Signaling, 1:1,000) or anti-mouse IgG HRP-linked (Cell Signaling, 1:1,000) with the SuperSignal^TM^ West Femto Chemiluminescent substrate (Thermo Fisher Scientific).

### Transcriptome sequencing

After RiboGreen quantification and quality control by Agilent BioAnalyzer, 0.5-1ng total RNA with RNA integrity numbers ranging from 6.1 to 10 underwent amplification using the SMART-Seq v4 Ultra Low Input RNA Kit (Clonetech catalog # 63488), with 12 cycles of amplification. Subsequently, 1.6-10ng of amplified cDNA was used to prepare libraries with the KAPA Hyper Prep Kit (Kapa Biosystems KK8504) using 8 cycles of PCR. Samples were barcoded and run on a HiSeq 4000 or NovaSeq 6000 in a PE50 run, using the HiSeq 3000/4000 SBS Kit or NovaSeq 6000 S1 Reagent Kit (100 Cycles) (Illumina). An average of 56 million paired reads were generated per sample and the percent of mRNA bases per sample ranged from 48% to 76%.

### RNA-Seq 1 day post transplantation

One day after the intracranial injection of mDA neurons sorted by NURR1-GFP+ signal, the mice were euthanized, and the injection site was grossly dissected and processed for papain dissociation (Worthington). Dissociated xenograft sample along with in vitro cultured neurons (day 1 *in vitro*) are simultaneously subject to FACS for re-isolating dopamine neurons based on the endogenous reporter signal. Total RNA was extracted in TRIzol (Invitrogen) according to the manufacturer’s instructions. RNAseq libraries of polyadenylated RNA were prepared using the TruSeq Stranded mRNA Library Prep Kit (Illumina) according to the manufacturer’s instructions and sequenced on an Illumina NextSeq 500 platform. The resulting single end reads were checked for quality (FastQC v0.11.5) and processed RNAseq libraries of polyadenylated RNA were prepared using the TruSeq Stranded mRNA Library Prep Kit (Illumina) according to the manufacturer’s instructions and sequenced on an Illumina NextSeq 500 platform. The resultant filtered reads were mapped to human reference genome hg19 using STAR aligner (version 2.5.0a). The expression data at the coding sequence were quantified using the GENCODE version 38 transcriptome (Frankish et al., 2019) and HTSeq version 1.99.2 (Anders et al., 2015). Differential expression analysis was done with the negative binomial statistical model using DESeq2 version 1.30.1 (Love et al., 2014). Enrichment analysis with the various reference data was done using clusterProfiler version 4.2.1 (Wu et al., 2021; Yu et al., 2012).

### Single-cell transcriptome sequencing

Single cell suspensions were stained with Trypan blue, and the Countess II Automated Cell Counter (ThermoFisher) was used to assess both cell number and viability. Following QC, the single cell suspension was loaded onto Chromium Chip B (10X Genomics PN 2000060) and GEM generation, cDNA synthesis, cDNA amplification, and library preparation of 3,900-5,300 cells proceeded using the Chromium Single Cell 3’ Reagent Kit v3 (10X Genomics PN 1000075) according to the manufacturer’s protocol. cDNA amplification included 12 cycles and 88-99ng of the material was used to prepare sequencing libraries with 12 cycles of PCR. Indexed libraries were pooled equimolar and sequenced on a NovaSeq 6000 in a PE28/91 paired end run using the NovaSeq 6000 S1 Reagent Kit (100 cycles) (Illumina). An average of 94 million paired reads was generated per sample.

### Single cell analysis

The samples underwent 10X chromium Single Cell 3’ v3 processing. The reads were aligned to human GRCh38 (GENCODE v32/Ensembl 98) using Cell Ranger 5.0.0. The resulting filtered count matrix was further filtered for cells with i) minimum 1000 UMI counts, ii) 500 ≤ gene counts ≤ 7000, iii) and mitochondrial gene percentage of less than 25%. Normalization by deconvolution in scran version 1.22.1 was performed and the signal from the gene expression related to the cell cycle was regressed out as directed by Seurat version 4.1. The default 2000 highly variable genes were selected, and the first 50 principal components were extracted from the cell cycle-regressed matrix. Subsequently, the shared nearest neighbors were calculated from the principal components using buildSNNGraph of R software scran using the k parameter of 40. Seven clusters were identified and using the walktrap algorithm, with the function cluster_walktrap of R implementation of the igraph package version 1.3.5. The uniform manifold approximation and projection (UMAP) was performed. Differential gene expression was performed via the Seurat package using MAST. Cluster annotation was performed via clusterProfiler package version 4.2.2, and differential expression visualization using EnhancedVolcano version 1.12.0.

GEO accession number: data submitted and accession number pending.

### Quantification and Statistical analysis

N=3 independent biological replicates were used for all experiments unless otherwise indicated. n.s. indicates a non-significant difference. *P*-values were calculated by unpaired two-tailed Student’s t-test unless otherwise indicated. \**p*<0.05, \*\**p*<0.01 and \*\*\**p*<0.001.

## CITED REFERENCES

Anders, S., Pyl, P.T., and Huber, W. (2015). HTSeq--a Python framework to work with high-throughput sequencing data. Bioinformatics 31, 166–169.

Barker, R.A., Parmar, M., Studer, L., and Takahashi, J. (2017). Human Trials of Stem Cell-Derived Dopamine Neurons for Parkinson’s Disease: Dawn of a New Era. Cell Stem Cell 21, 569–573.

Bloem, B.R., Okun, M.S., and Klein, C. (2021). Parkinson’s disease. Lancet 397, 2284–2303.

Chen, S., Han, Y., Yang, L., Kim, T., Nair, M., Harschnitz, O., Wang, P., Zhu, J., Koo, S.Y., Tang, X., et al. (2021). SARS-CoV-2 Infection Causes Dopaminergic Neuron Senescence. Res Sq.

Chen, S., Sanjana, N.E., Zheng, K., Shalem, O., Lee, K., Shi, X., Scott, D.A., Song, J., Pan, J.Q., Weissleder, R., et al. (2015). Genome-wide CRISPR screen in a mouse model of tumor growth and metastasis. Cell 160, 1246–1260.

Chong, J.J., Yang, X., Don, C.W., Minami, E., Liu, Y.W., Weyers, J.J., Mahoney, W.M., Van Biber, B., Cook, S.M., Palpant, N.J., et al. (2014). Human embryonic-stem-cell-derived cardiomyocytes regenerate non-human primate hearts. Nature 510, 273–277.

Chou, J., Greig, N.H., Reiner, D., Hoffer, B.J., and Wang, Y. (2011). Enhanced survival of dopaminergic neuronal transplants in hemiparkinsonian rats by the p53 inactivator PFT-alpha. Cell Transplant 20, 1351–1359.

Courtois, E.T., Castillo, C.G., Seiz, E.G., Ramos, M., Bueno, C., Liste, I., and Martinez-Serrano, A. (2010). In vitro and in vivo enhanced generation of human A9 dopamine neurons from neural stem cells by Bcl-XL. J Biol Chem 285, 9881–9897.

Doi, D., Magotani, H., Kikuchi, T., Ikeda, M., Hiramatsu, S., Yoshida, K., Amano, N., Nomura, M., Umekage, M., Morizane, A., et al. (2020). Pre-clinical study of induced pluripotent stem cell-derived dopaminergic progenitor cells for Parkinson’s disease. Nat Commun 11, 3369.

Doi, D., Samata, B., Katsukawa, M., Kikuchi, T., Morizane, A., Ono, Y., Sekiguchi, K., Nakagawa, M., Parmar, M., and Takahashi, J. (2014). Isolation of human induced pluripotent stem cell-derived dopaminergic progenitors by cell sorting for successful transplantation. Stem Cell Reports 2, 337–350.

Dorsey, E.R., Sherer, T., Okun, M.S., and Bloem, B.R. (2018). The Emerging Evidence of the Parkinson Pandemic. J Parkinsons Dis 8, S3–S8.

Duan, W.M., Widner, H., and Brundin, P. (1995). Temporal pattern of host responses against intrastriatal grafts of syngeneic, allogeneic or xenogeneic embryonic neuronal tissue in rats. Exp Brain Res 104, 227–242.

Frankish, A., Diekhans, M., Ferreira, A.M., Johnson, R., Jungreis, I., Loveland, J., Mudge, J.M., Sisu, C., Wright, J., Armstrong, J., et al. (2019). GENCODE reference annotation for the human and mouse genomes. Nucleic Acids Res 47, D766–D773.

Freed, C.R., Greene, P.E., Breeze, R.E., Tsai, W.Y., DuMouchel, W., Kao, R., Dillon, S., Winfield, H., Culver, S., Trojanowski, J.Q., et al. (2001). Transplantation of embryonic dopamine neurons for severe Parkinson’s disease. N Engl J Med 344, 710–719.

Ganat, Y.M., Calder, E.L., Kriks, S., Nelander, J., Tu, E.Y., Jia, F., Battista, D., Harrison, N., Parmar, M., Tomishima, M.J., et al. (2012). Identification of embryonic stem cell-derived midbrain dopaminergic neurons for engraftment. J Clin Invest 122, 2928–2939.

Gedye, C.A., Hussain, A., Paterson, J., Smrke, A., Saini, H., Sirskyj, D., Pereira, K., Lobo, N., Stewart, J., Go, C., et al. (2014). Cell surface profiling using high-throughput flow cytometry: a platform for biomarker discovery and analysis of cellular heterogeneity. PLoS One 9, e105602.

Gonzalez, F., Zhu, Z., Shi, Z.D., Lelli, K., Verma, N., Li, Q.V., and Huangfu, D. (2014). An iCRISPR platform for rapid, multiplexable, and inducible genome editing in human pluripotent stem cells. Cell Stem Cell 15, 215–226.

Hafner, A., Bulyk, M.L., Jambhekar, A., and Lahav, G. (2019). The multiple mechanisms that regulate p53 activity and cell fate. Nat Rev Mol Cell Biol 20, 199–210.

Kamath, T., Abdulraouf, A., Burris, S., Gazestani, V., Nadaf, N., Vanderburg, C., and Macosko, E.Z. (2021). A molecular census of midbrain dopaminergic neurons in Parkinson’s disease. bioRxiv, 2021.2006.2016.448661.

Kamath, T., Abdulraouf, A., Burris, S.J., Langlieb, J., Gazestani, V., Nadaf, N.M., Balderrama, K., Vanderburg, C., and Macosko, E.Z. (2022). Single-cell genomic profiling of human dopamine neurons identifies a population that selectively degenerates in Parkinson’s disease. Nat Neurosci 25, 588–595.

Kefalopoulou, Z., Politis, M., Piccini, P., Mencacci, N., Bhatia, K., Jahanshahi, M., Widner, H., Rehncrona, S., Brundin, P., Bjorklund, A., et al. (2014). Long-term clinical outcome of fetal cell transplantation for Parkinson disease: two case reports. JAMA Neurol 71, 83–87.

Kikuchi, T., Morizane, A., Doi, D., Magotani, H., Onoe, H., Hayashi, T., Mizuma, H., Takara, S., Takahashi, R., Inoue, H., et al. (2017). Human iPS cell-derived dopaminergic neurons function in a primate Parkinson’s disease model. Nature 548, 592–596.

Kim, T.W., Koo, S.Y., and Studer, L. (2020). Pluripotent Stem Cell Therapies for Parkinson Disease: Present Challenges and Future Opportunities. Front Cell Dev Biol 8, 729.

Kim, T.W., Piao, J., Koo, S.Y., Kriks, S., Chung, S.Y., Betel, D., Socci, N.D., Choi, S.J., Zabierowski, S., Dubose, B.N., et al. (2021). Biphasic Activation of WNT Signaling Facilitates the Derivation of Midbrain Dopamine Neurons from hESCs for Translational Use. Cell Stem Cell 28, 343–355 e345.

Kirkeby, A., Nolbrant, S., Tiklova, K., Heuer, A., Kee, N., Cardoso, T., Ottosson, D.R., Lelos, M.J., Rifes, P., Dunnett, S.B., et al. (2017). Predictive Markers Guide Differentiation to Improve Graft Outcome in Clinical Translation of hESC-Based Therapy for Parkinson’s Disease. Cell Stem Cell 20, 135–148.

Kolde, R. (2018). Package ‘pheatmap’. In R Package (https://cran.microsoft.com/snapshot/2018-06-22/web/packages/pheatmap/pheatmap.pdf).

Kriks, S., Shim, J.W., Piao, J., Ganat, Y.M., Wakeman, D.R., Xie, Z., Carrillo-Reid, L., Auyeung, G., Antonacci, C., Buch, A., et al. (2011). Dopamine neurons derived from human ES cells efficiently engraft in animal models of Parkinson’s disease. Nature 480, 547–551.

La Manno, G., Gyllborg, D., Codeluppi, S., Nishimura, K., Salto, C., Zeisel, A., Borm, L.E., Stott, S.R.W., Toledo, E.M., Villaescusa, J.C., et al. (2016). Molecular Diversity of Midbrain Development in Mouse, Human, and Stem Cells. Cell 167, 566–580 e519.

Lehnen, D., Barral, S., Cardoso, T., Grealish, S., Heuer, A., Smiyakin, A., Kirkeby, A., Kollet, J., Cremer, H., Parmar, M., et al. (2017). IAP-Based Cell Sorting Results in Homogeneous Transplantable Dopaminergic Precursor Cells Derived from Human Pluripotent Stem Cells. Stem Cell Reports 9, 1207–1220.

Li, W., Englund, E., Widner, H., Mattsson, B., van Westen, D., Latt, J., Rehncrona, S., Brundin, P., Bjorklund, A., Lindvall, O., et al. (2016). Extensive graft-derived dopaminergic innervation is maintained 24 years after transplantation in the degenerating parkinsonian brain. Proc Natl Acad Sci U S A 113, 6544–6549.

Liddelow, S.A., Guttenplan, K.A., Clarke, L.E., Bennett, F.C., Bohlen, C.J., Schirmer, L., Bennett, M.L., Munch, A.E., Chung, W.S., Peterson, T.C., et al. (2017). Neurotoxic reactive astrocytes are induced by activated microglia. Nature 541, 481–487.

Liu, G., Yu, J., Ding, J., Xie, C., Sun, L., Rudenko, I., Zheng, W., Sastry, N., Luo, J., Rudow, G., et al. (2014). Aldehyde dehydrogenase 1 defines and protects a nigrostriatal dopaminergic neuron subpopulation. J Clin Invest 124, 3032–3046.

Love, M.I., Huber, W., and Anders, S. (2014). Moderated estimation of fold change and dispersion for RNA-seq data with DESeq2. Genome Biol 15, 550.

Maor-Nof, M., Shipony, Z., Lopez-Gonzalez, R., Nakayama, L., Zhang, Y.J., Couthouis, J., Blum, J.A., Castruita, P.A., Linares, G.R., Ruan, K., et al. (2021). p53 is a central regulator driving neurodegeneration caused by C9orf72 poly(PR). Cell 184, 689–708 e620.

McCarthy, D.J., Chen, Y., and Smyth, G.K. (2012). Differential expression analysis of multifactor RNA-Seq experiments with respect to biological variation. Nucleic Acids Res 40, 4288–4297.

McCoy, M.K., and Tansey, M.G. (2008). TNF signaling inhibition in the CNS: implications for normal brain function and neurodegenerative disease. J Neuroinflammation 5, 45.

Michels, B.E., Mosa, M.H., Streibl, B.I., Zhan, T., Menche, C., Abou-El-Ardat, K., Darvishi, T., Czlonka, E., Wagner, S., Winter, J., et al. (2020). Pooled In Vitro and In Vivo CRISPR-Cas9 Screening Identifies Tumor Suppressors in Human Colon Organoids. Cell Stem Cell 26, 782–792 e787.

Monzon-Sandoval, J., Poggiolini, I., Ilmer, T., Wade-Martins, R., Webber, C., and Parkkinen, L. (2020). Human-Specific Transcriptome of Ventral and Dorsal Midbrain Dopamine Neurons. Ann Neurol 87, 853–868.

Nguyen, H.X., Hooshmand, M.J., Saiwai, H., Maddox, J., Salehi, A., Lakatos, A., Nishi, R.A., Salazar, D., Uchida, N., and Anderson, A.J. (2017). Systemic Neutrophil Depletion Modulates the Migration and Fate of Transplanted Human Neural Stem Cells to Rescue Functional Repair. J Neurosci 37, 9269–9287.

Olanow, C.W., Goetz, C.G., Kordower, J.H., Stoessl, A.J., Sossi, V., Brin, M.F., Shannon, K.M., Nauert, G.M., Perl, D.P., Godbold, J., et al. (2003). A double-blind controlled trial of bilateral fetal nigral transplantation in Parkinson’s disease. Ann Neurol 54, 403–414.

Panman, L., Papathanou, M., Laguna, A., Oosterveen, T., Volakakis, N., Acampora, D., Kurtsdotter, I., Yoshitake, T., Kehr, J., Joodmardi, E., et al. (2014). Sox6 and Otx2 control the specification of substantia nigra and ventral tegmental area dopamine neurons. Cell Rep 8, 1018–1025.

Parmar, M., Grealish, S., and Henchcliffe, C. (2020). The future of stem cell therapies for Parkinson disease. Nat Rev Neurosci 21, 103–115.

Pasca, A.M., Park, J.Y., Shin, H.W., Qi, Q., Revah, O., Krasnoff, R., O’Hara, R., Willsey, A.J., Palmer, T.D., and Pasca, S.P. (2019). Human 3D cellular model of hypoxic brain injury of prematurity. Nat Med 25, 784–791.

Pereira Luppi, M., Azcorra, M., Caronia-Brown, G., Poulin, J.F., Gaertner, Z., Gatica, S., Moreno-Ramos, O.A., Nouri, N., Dubois, M., Ma, Y.C., et al. (2021). Sox6 expression distinguishes dorsally and ventrally biased dopamine neurons in the substantia nigra with distinctive properties and embryonic origins. Cell Rep 37, 109975.

Perez, A.R., Pritykin, Y., Vidigal, J.A., Chhangawala, S., Zamparo, L., Leslie, C.S., and Ventura, A. (2017). GuideScan software for improved single and paired CRISPR guide RNA design. Nat Biotechnol 35, 347–349.

Piao, J., Zabierowski, S., Dubose, B.N., Hill, E.J., Navare, M., Claros, N., Rosen, S., Ramnarine, K., Horn, C., Fredrickson, C., et al. (2021). Preclinical Efficacy and Safety of a Human Embryonic Stem Cell-Derived Midbrain Dopamine Progenitor Product, MSK-DA01. Cell Stem Cell *28*, 217-229 e217.

Poewe, W., Seppi, K., Tanner, C.M., Halliday, G.M., Brundin, P., Volkmann, J., Schrag, A.E., and Lang, A.E. (2017). Parkinson disease. Nat Rev Dis Primers 3, 17013.

Politis, M., Wu, K., Loane, C., Quinn, N.P., Brooks, D.J., Rehncrona, S., Bjorklund, A., Lindvall, O., and Piccini, P. (2010). Serotonergic neurons mediate dyskinesia side effects in Parkinson’s patients with neural transplants. Sci Transl Med 2, 38ra46.

Riessland, M., Kolisnyk, B., Kim, T.W., Cheng, J., Ni, J., Pearson, J.A., Park, E.J., Dam, K., Acehan, D., Ramos-Espiritu, L.S., et al. (2019). Loss of SATB1 Induces p21-Dependent Cellular Senescence in Post-mitotic Dopaminergic Neurons. Cell Stem Cell 25, 514–530 e518.

Robinson, M.D., McCarthy, D.J., and Smyth, G.K. (2010). edgeR: a Bioconductor package for differential expression analysis of digital gene expression data. Bioinformatics 26, 139–140.

Samata, B., Doi, D., Nishimura, K., Kikuchi, T., Watanabe, A., Sakamoto, Y., Kakuta, J., Ono, Y., and Takahashi, J. (2016). Purification of functional human ES and iPSC-derived midbrain dopaminergic progenitors using LRTM1. Nat Commun 7, 13097.

Savitt, J.M., Jang, S.S., Mu, W., Dawson, V.L., and Dawson, T.M. (2005). Bcl-x is required for proper development of the mouse substantia nigra. J Neurosci 25, 6721–6728.

Schwartzentruber, J., Foskolou, S., Kilpinen, H., Rodrigues, J., Alasoo, K., Knights, A.J., Patel, M., Goncalves, A., Ferreira, R., Benn, C.L., et al. (2018). Molecular and functional variation in iPSC-derived sensory neurons. Nat Genet 50, 54–61.

Schweitzer, J.S., Song, B., Herrington, T.M., Park, T.Y., Lee, N., Ko, S., Jeon, J., Cha, Y., Kim, K., Li, Q., et al. (2020). Personalized iPSC-Derived Dopamine Progenitor Cells for Parkinson’s Disease. N Engl J Med 382, 1926–1932.

Shalem, O., Sanjana, N.E., and Zhang, F. (2015). High-throughput functional genomics using CRISPR-Cas9. Nat Rev Genet 16, 299–311.

Shim, J.W., Koh, H.C., Chang, M.Y., Roh, E., Choi, C.Y., Oh, Y.J., Son, H., Lee, Y.S., Studer, L., and Lee, S.H. (2004). Enhanced in vitro midbrain dopamine neuron differentiation, dopaminergic function, neurite outgrowth, and 1-methyl-4-phenylpyridium resistance in mouse embryonic stem cells overexpressing Bcl-XL. J Neurosci 24, 843–852.

Spangenberg, E.E., Lee, R.J., Najafi, A.R., Rice, R.A., Elmore, M.R., Blurton-Jones, M., West, B.L., and Green, K.N. (2016). Eliminating microglia in Alzheimer’s mice prevents neuronal loss without modulating amyloid-beta pathology. Brain 139, 1265–1281.

Sun, C., Kannan, S., Choi, I.Y., Lim, H., Zhang, H., Chen, G.S., Zhang, N., Park, S.H., Serra, C., Iyer, S.R., et al. (2022). Human pluripotent stem cell-derived myogenic progenitors undergo maturation to quiescent satellite cells upon engraftment. Cell Stem Cell 29, 610–619 e615.

Surmeier, D.J., Obeso, J.A., and Halliday, G.M. (2017). Selective neuronal vulnerability in Parkinson disease. Nat Rev Neurosci 18, 101–113.

Tao, Y., Vermilyea, S.C., Zammit, M., Lu, J., Olsen, M., Metzger, J.M., Yao, L., Chen, Y., Phillips, S., Holden, J.E., et al. (2021). Autologous transplant therapy alleviates motor and depressive behaviors in parkinsonian monkeys. Nat Med 27, 632–639.

Tiklova, K., Nolbrant, S., Fiorenzano, A., Bjorklund, A.K., Sharma, Y., Heuer, A., Gillberg, L., Hoban, D.B., Cardoso, T., Adler, A.F., et al. (2020). Single cell transcriptomics identifies stem cell-derived graft composition in a model of Parkinson’s disease. Nat Commun 11, 2434.

Tuttolomondo, A., Pecoraro, R., and Pinto, A. (2014). Studies of selective TNF inhibitors in the treatment of brain injury from stroke and trauma: a review of the evidence to date. Drug Des Devel Ther 8, 2221–2238.

Villaescusa, J.C., Li, B., Toledo, E.M., Rivetti di Val Cervo, P., Yang, S., Stott, S.R., Kaiser, K., Islam, S., Gyllborg, D., Laguna-Goya, R., et al. (2016). A PBX1 transcriptional network controls dopaminergic neuron development and is impaired in Parkinson’s disease. EMBO J 35, 1963–1978.

Volpato, V., Smith, J., Sandor, C., Ried, J.S., Baud, A., Handel, A., Newey, S.E., Wessely, F., Attar, M., Whiteley, E., et al. (2018). Reproducibility of Molecular Phenotypes after Long-Term Differentiation to Human iPSC-Derived Neurons: A Multi-Site Omics Study. Stem Cell Reports 11, 897–911.

Wang, E., Zhou, H., Nadorp, B., Cayanan, G., Chen, X., Yeaton, A.H., Nomikou, S., Witkowski, M.T., Narang, S., Kloetgen, A., et al. (2021a). Surface antigen-guided CRISPR screens identify regulators of myeloid leukemia differentiation. Cell Stem Cell 28, 718–731 e716.

Wang, X., Tokheim, C., Gu, S.S., Wang, B., Tang, Q., Li, Y., Traugh, N., Zeng, Z., Zhang, Y., Li, Z., et al. (2021b). In vivo CRISPR screens identify the E3 ligase Cop1 as a modulator of macrophage infiltration and cancer immunotherapy target. Cell 184, 5357–5374 e5322.

Winkler, C., Kirik, D., and Bjorklund, A. (2005). Cell transplantation in Parkinson’s disease: how can we make it work? Trends Neurosci 28, 86–92.

Wu, T., Hu, E., Xu, S., Chen, M., Guo, P., Dai, Z., Feng, T., Zhou, L., Tang, W., Zhan, L., et al. (2021). clusterProfiler 4.0: A universal enrichment tool for interpreting omics data. Innovation (N Y) 2, 100141.

Yu, G., Wang, L.G., Han, Y., and He, Q.Y. (2012). clusterProfiler: an R package for comparing biological themes among gene clusters. OMICS 16, 284–287.

