## Supplemental Figures for "TNF-NFkB-p53 axis restricts *in vivo* survival of hPSC-derived dopamine neuron"

### Supplementary Figure 1

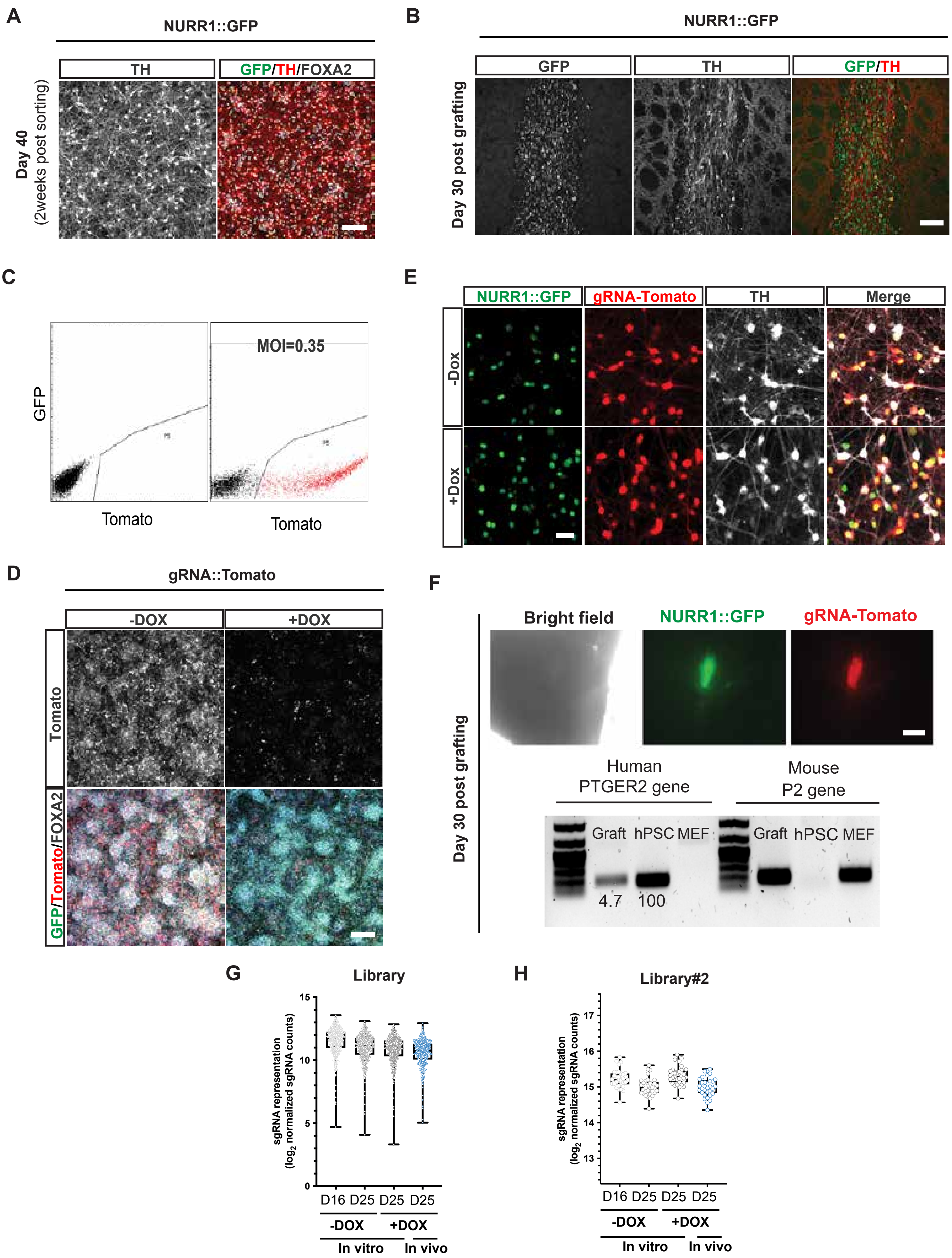

### Supplementary Figure 2

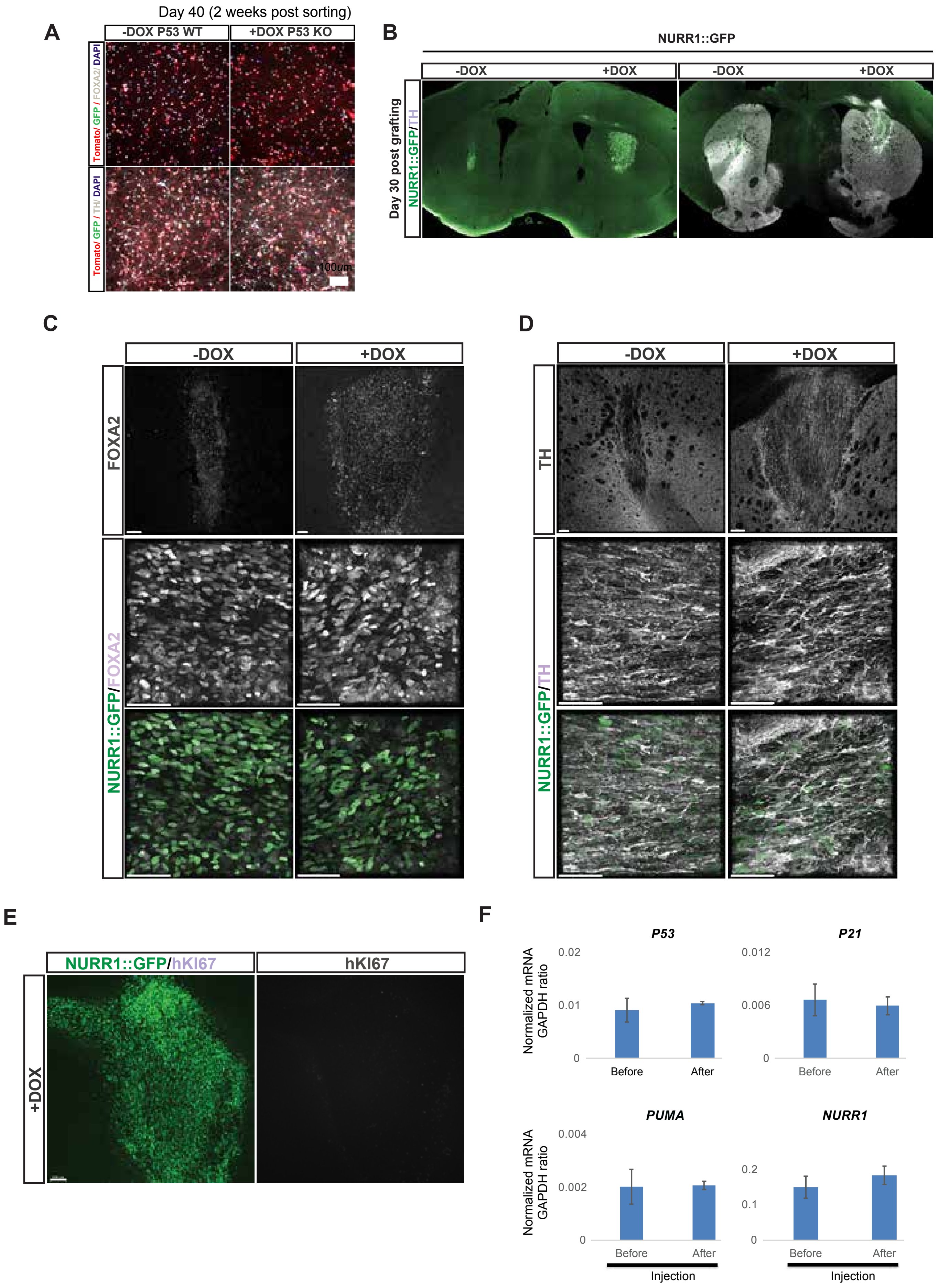

Supplementary Figure 3

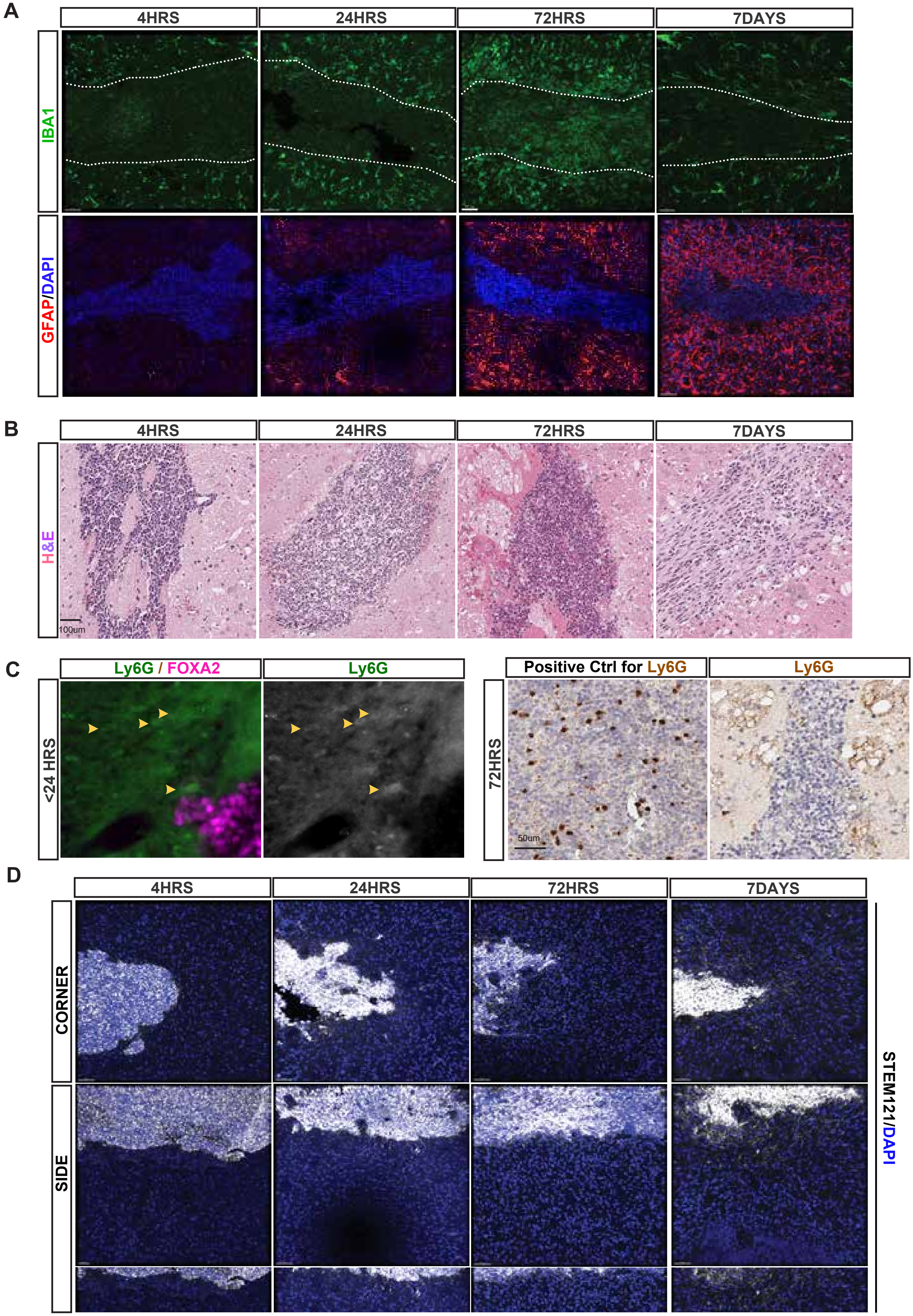

### Supplementary Figure 4

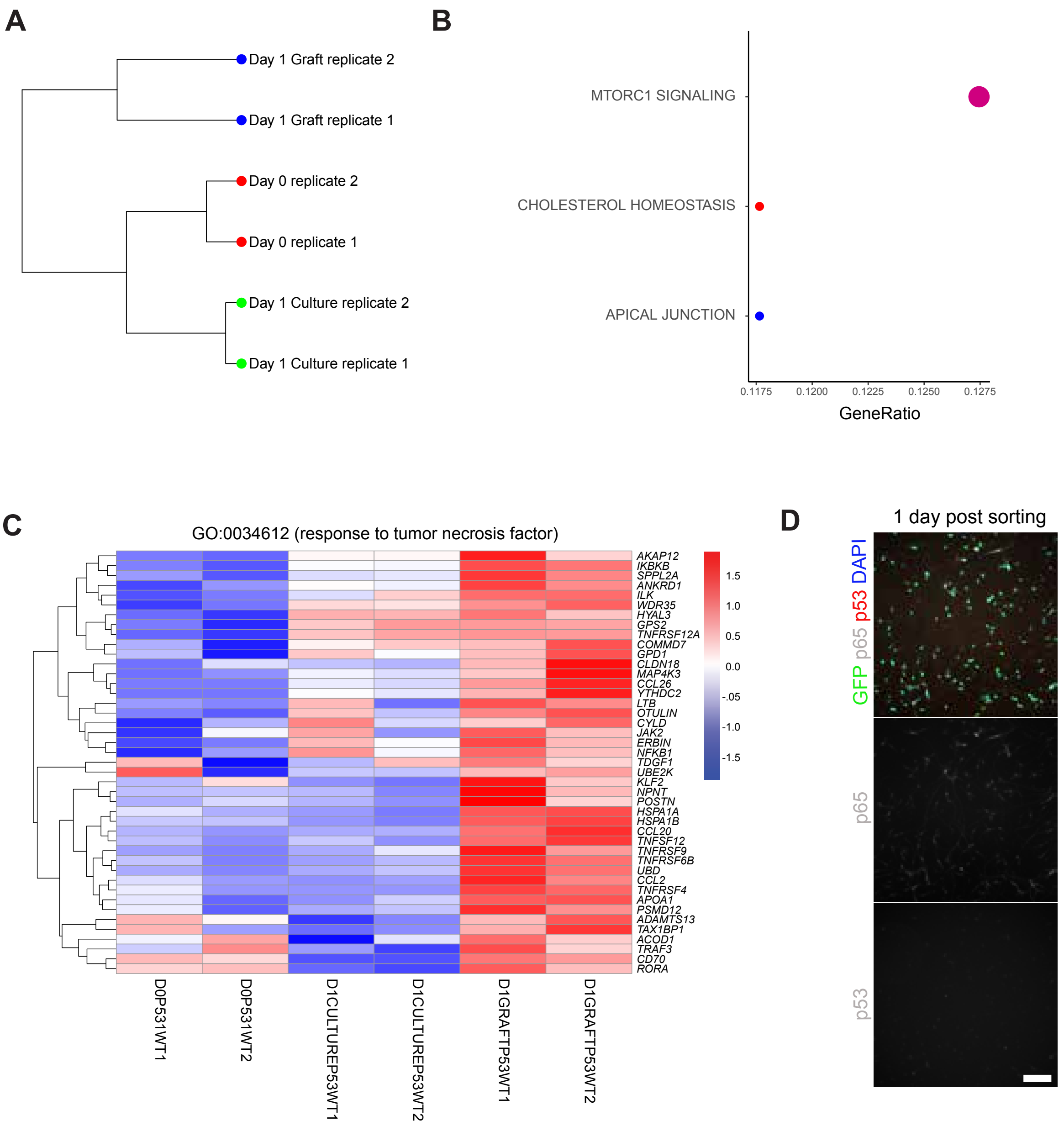

### Supplementary Figure 5

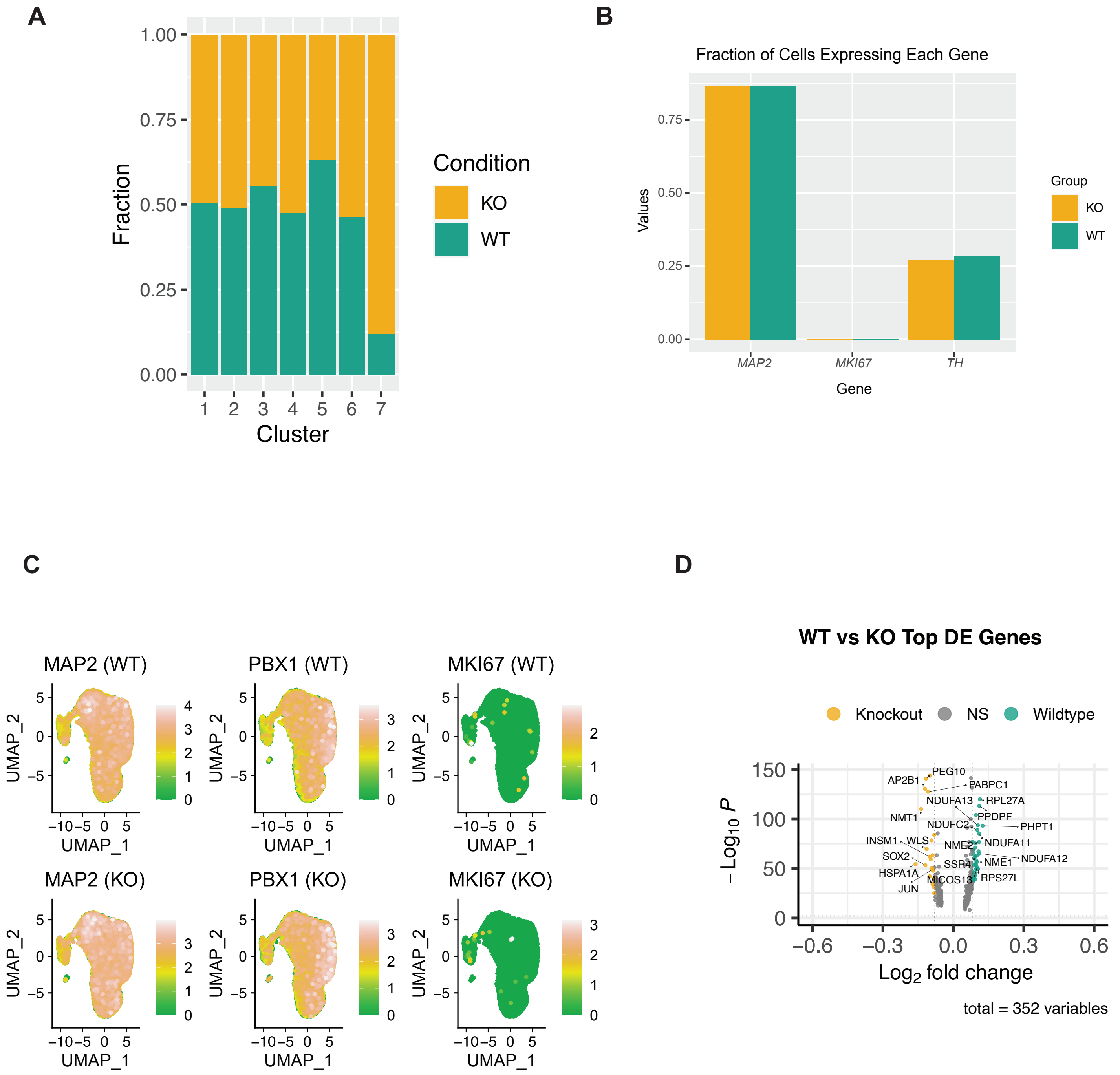

#### Supplementary Figure 6

**A**

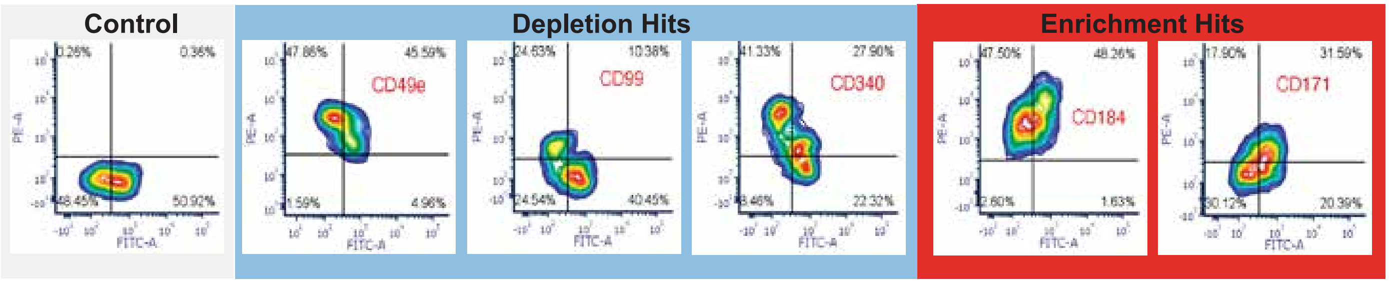


**B**

**30 days post grafting**

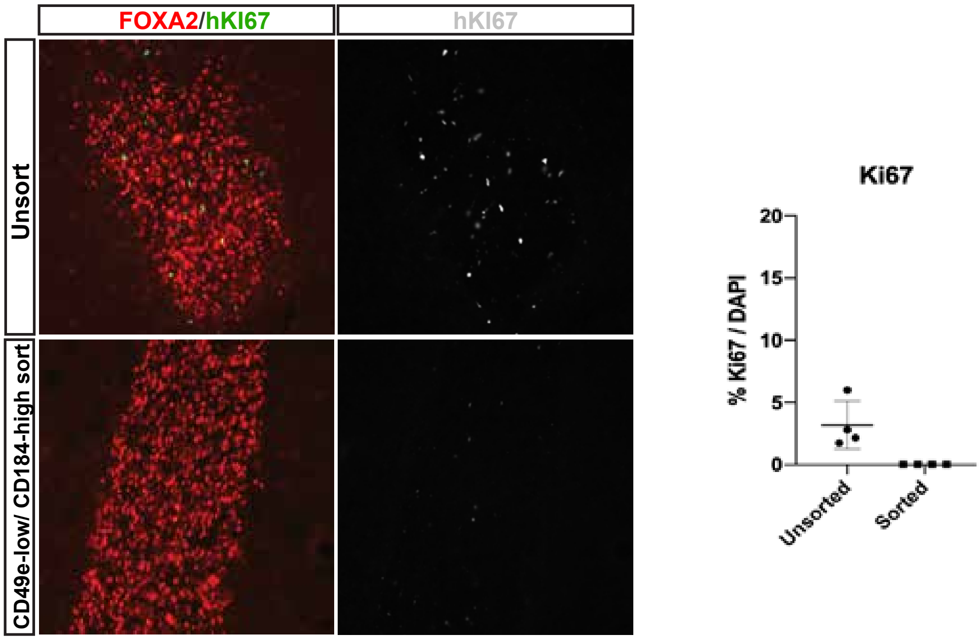

### Supplementary Figure 7

A

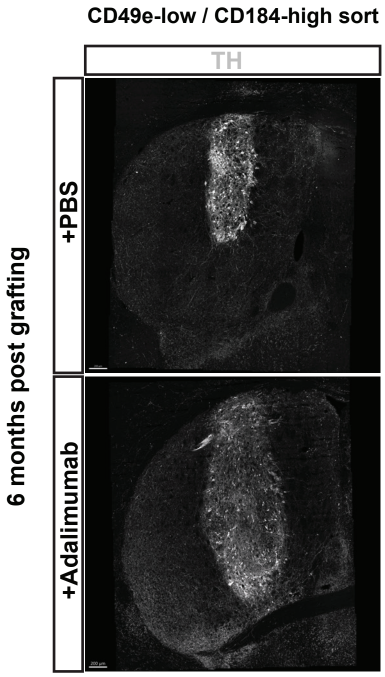


B

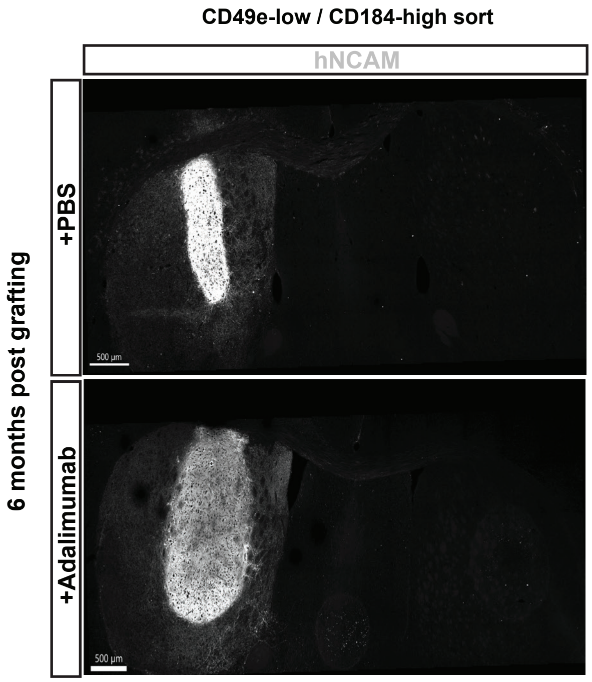


C

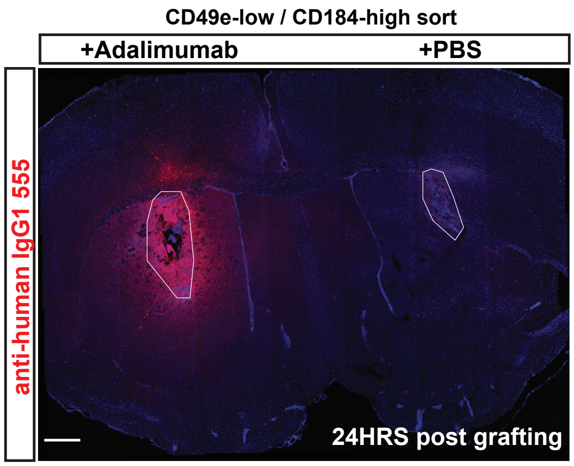
